# Defective Desmosomal Adhesion Causes Arrhythmogenic Cardiomyopathy by involving an Integrin-αVβ6/TGF-β Signaling Cascade

**DOI:** 10.1101/2021.09.02.458734

**Authors:** Camilla Schinner, Henriette Franz, Aude Zimmermann, Marie-Therès Wanuske, Florian Geier, Pawel Pelczar, Vera Lorenz, Lifen Xu, Chiara Stüdle, Piotr I Maly, Silke Kauferstein, Britt Maria Beckmann, Gabriela M Kuster, Volker Spindler

## Abstract

**Background:** Arrhythmogenic Cardiomyopathy (ACM) is characterized by progressive loss of cardiomyocytes with fibrofatty replacement, systolic dysfunction and life-threatening arrhythmias. A substantial proportion of ACM is caused by mutations in genes of the desmosomal cell-cell adhesion complex, but the underlying mechanisms are not well understood. So far, treatment options are only symptomatic. Here, we investigate the relevance of defective desmosomal adhesion for ACM development and progression.

**Methods:** We mutated the binding site of desmoglein-2 (DSG2), a crucial desmosomal adhesion molecule in cardiomyocytes. This DSG2-W2A mutation abrogates the tryptophan swap, a central interaction mechanism of DSG2 based on structural data. Impaired adhesive function of DSG2-W2A was confirmed by cell-cell dissociation assays and force spectroscopy measurements by atomic force microscopy. We next generated a DSG2-W2A knock-in mouse model, which was analyzed by echocardiography and histological and bio-molecular techniques including RNA sequencing, transmission electron and super-resolution microscopy. The results were compared to ACM patient samples and their relevance was confirmed in cardiac slice cultures.

**Results:** The DSG2-W2A mutation induced impaired binding and desmosomal adhesion dysfunction on cellular and molecular level. Mice bearing this mutation develop a severe cardiac phenotype recalling the characteristics of ACM, including cardiac fibrosis, impaired systolic function and arrhythmia. A comparison of the transcriptome of mutant mice with ACM patient data suggested deregulated integrin-αVβ6 and subsequent TGF-β signaling as driver of cardiac fibrosis. Accordingly, blocking antibodies targeting integrin-αVβ6 or inhibition of TGF-β receptor signaling both led to reduced expression of pro-fibrotic markers in cardiac slice cultures.

**Conclusions:** Here, we show that disruption of desmosomal adhesion is sufficient to induce ACM, which confirms the dysfunctional adhesion hypothesis. Mechanistically, deregulation of integrin-αVβ6 signaling was identified as a central step towards fibrosis. This highlights the value of this model to discern mechanisms of cardiac fibrosis and to identify and test novel treatment options for ACM.

## Introduction

Patients suffering from Arrhythmogenic Cardiomyopathy (ACM) are presenting with impaired cardiac function and ventricular arrhythmia up to sudden cardiac death.^1, 2^ This inherited disease occurs with a prevalence of 1:1’000 – 1:5’000 and becomes typically evident in adults between 20 and 40 years of age.^3^ Structurally, ACM is characterized by progressive loss of cardiomyocytes, cardiac fibrosis with fibrofatty tissue replacement and ventricular dilatation. In the majority of cases, mutations in components of the desmosomal complex are causative for ACM.^1, 2, 4^

Desmosomes are intercellular junctions which provide strong cell-cell adhesion in mechanically challenged tissues such as the myocardium or epithelia. Within desmosomes, desmosomal cadherins, namely desmogleins (DSGs, isoform 1-4) and desmocollins (DSCs, isoform 1-3), protrude into the intercellular space to interact with the corresponding molecules from the opposing cell. The cytoplasmic tail of these transmembrane adhesion molecules is anchored to the intermediate filament system via the plaque proteins plakoglobin (PG), plakophillins (PKPs, isoform 1-3), and desmoplakin (DSP). This setup confers remarkable resilience against mechanical forces.^5^ In cardiomyocytes, only DSG2 and DSC2 are expressed and are linked to the desmin intermediate filament system via PG, PKP2 and DSP.^6^ These desmosomal components are an integral part of the intercalated disc (ICD) where they form a functional unit with molecules of adherens junctions, gap junctions, and others such as sodium channels.^7–9^ The ICD is essential to provide mechanical and electrical coupling of adjacent cardiomyocytes, which is the basis for coordinated contraction and function of the entire heart.

In ACM, a variety of mutations in genes encoding for desmosomal molecules were described.^2^ Together with the finding of disrupted ICD ultrastructure,^10^ this formed the hypothesis of impaired cardiomyocyte cohesion as central step in ACM pathogenesis. In line with this, ACM seems to be aggravated by physical exercise and is one of the main causes of sudden cardiac death in athletes.^11, 12^ However, data on the role of cell-cell adhesion are conflicting and the impact of disrupted mechanical coupling for disease induction and progression is not finally resolved.^13–16^ Moreover, the mechanisms of fibrosis formation and arrhythmia as important hallmarks of the disease are not clear.^2, 17^ Thus, up to now, the therapeutic options for ACM are limited to symptomatic treatments like beta-adrenergic receptor blockers, restriction of physical exercise, implantation of a cardiac defibrillator, or heart transplantation as ultima ratio.^1^ This highlights the need for mouse models mimicking the ACM phenotype to identify new targeted treatment strategies to prevent disease development and progression. To this end, knock-out models for each desmosomal molecule profoundly increased the understanding of ACM patho-mechanisms. In addition, models overexpressing specific ACM patient mutations were generated.^2, 18^ However, protein deficiency restricted to cardiomyocytes as well as overexpression of mutated proteins under external promotors can only partially mimic human ACM.

In this study, we aim to investigate the role of disrupted desmosomal adhesion in ACM development by interfering with the binding mechanism of DSG2. Recently, the published crystal structure of DSG2 suggested the extracellular *trans*-interaction of desmosomal adhesion molecules to be mediated by a so-called tryptophane swap (Figure 1A).^19^ Here, a tryptophane residue at position 2 is inserting into a hydrophobic pocket of the first extracellular domain of the respective molecule from the opposing cell and vice versa. This hand shake-like exchange of tryptophane residues is proposed to mediate adhesion of DSGs and DSCs similar to the known binding mechanism of classical cadherins.^20^

**Figure 1.**
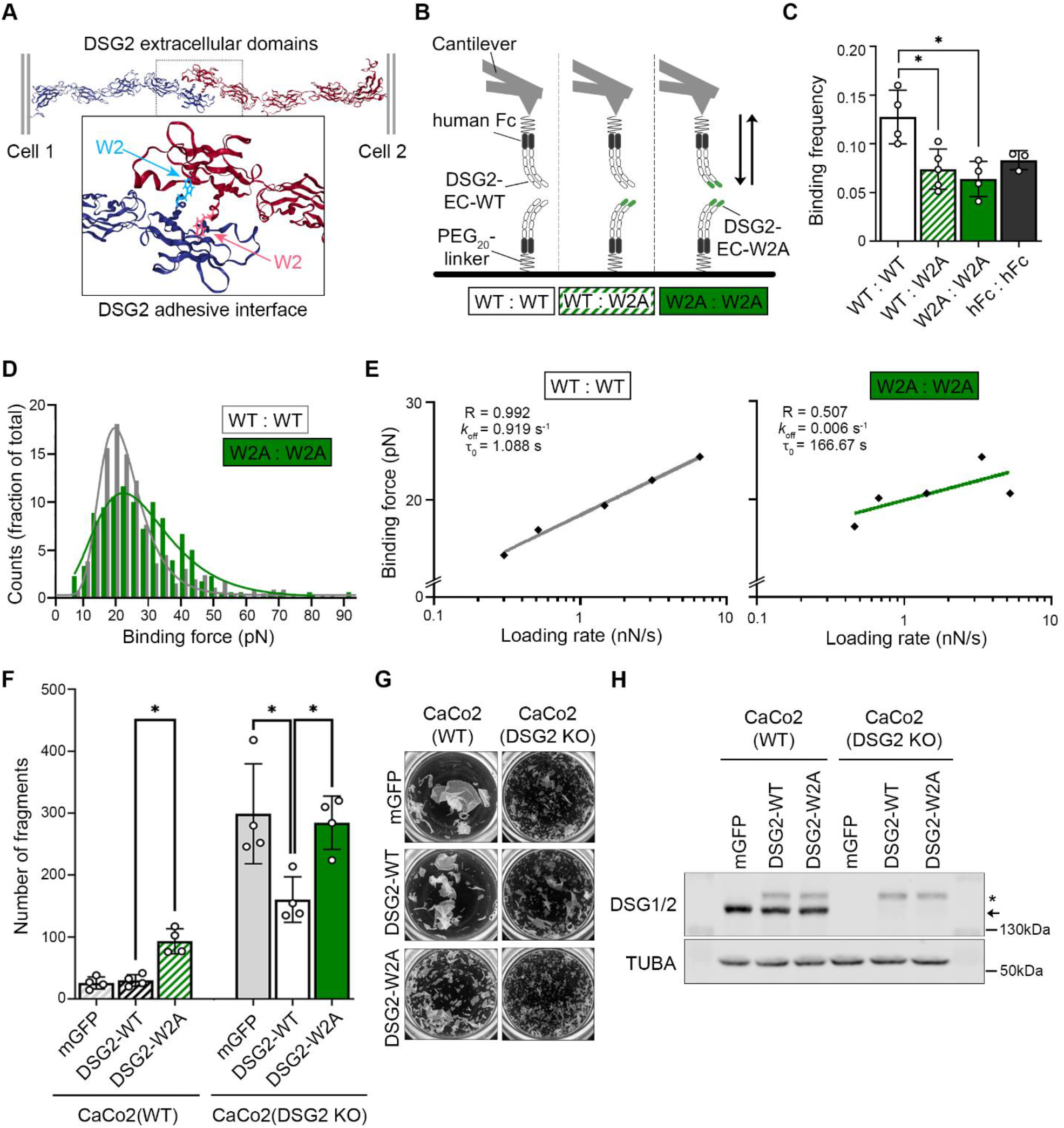
Desmoglein-2 interaction is mediated via tryptophane swap at position 2. (**A**) Predicted interaction model of desmoglein-2 (DSG2) extracellular domains via exchange of tryptophane residue at position 2 into a hydrophobic pocket of the opposing molecule. Cartoon 3D presentation of PDB entry 5ERD^19^, tryptophane-2 highlighted by ball and stick presentation. (**B**) Schematic of singe molecule force spectroscopy experiments. Recombinant extracellular domains (EC) of DSG2-WT or - W2A protein were coupled to a mica surface and AFM cantilever via a human Fc-tag (hFc) and a PEG_20_-linker and probed as indicated. (**C**) Interaction frequency of DSG2-W2A/DSG2-WT heterotypic and homotypic interactions at a pulling speed of 2 µm/s. Human Fc (hFc) served as control for unspecific binding. *P< 0.05, one-way ANOVA, Dunnett’s post hoc test. Each independent coating procedure is taken as biological replicate. (**D**) Histogram of binding forces distribution with peak fit at a pulling speed of 2 µm/s corresponding to data in **C**. (**E**) Determination of the bond half life time via Bell’s equation^21^ of mean loading rate and binding forces analysed from data of pulling speeds at 0.5, 1, 2, 5, 7.5 µm/s. Average of values from four independent coating procedures. R = R squared, *k*_off_ = off rate constant, τ_0_ = bond half lifetime under zero force. (**F**) Dissociation assays to determine cell-cell adhesion were performed in CaCo2 cells (WT or DSG2 KO background) expressing DSG2-WT-mGFP or DSG2-W2A-mGFP constructs. mGFP empty vector served as control. *P< 0.05, one-way ANOVA, Sidak’s post hoc test. (**G**) displays representative images of monolayer fragmentation from experiments in **F**. (**H**) Corresponding Western blot analysis confirmed effective expression of DSG2 constructs (*) vs. the endogenous protein (arrow) in CaCo2 cells. α-tubulin (TUBA) served as loading control.

To interfere with this mechanism, we inserted a mutation substituting the tryptophane-2 with an alanine (DSG2-W2A) and characterized its functional implication in vitro, in cultivated cells, and in a knock-in mouse model. Further, we applied this in vivo model to identify mechanisms underlying ACM aiming for new potential therapeutic approaches to treat ACM.

## Results

### DSG2-W2A mutation is abrogating desmosomal adhesion

To study the functional consequences of impaired DSG2 binding, we generated the W2A point mutation (DSG2-W2A) to abrogate the proposed interaction mechanism of the only desmoglein expressed in cardiomyocytes (Figure 1A). First, we investigated in a cell-free setup whether DSG2-W2A is indeed affecting the binding properties of DSG2 interaction by applying single molecule force spectroscopy experiments. Here, the extracellular domains of adhesion molecules are coupled to the tip of a sensitive atomic-force microscopy (AFM) probe and the surface of a mica sheet via a PEG linker. By measuring the deflection of the probe during repeated approach to and retraction from the surface, binding events can be quantitatively assessed. From these data, properties such as binding frequency, binding force, and bond half-life time can be determined. Constructs were generated for eukaryotic expression of wild type (WT) and mutant (W2A) DSG2 extracellular domains fused to a human Fc-fragment and tested for homo-(WT:WT, W2A:W2A) and heterotypic (WT:W2A) interaction properties (Figure 1B). Probing human Fc alone served as control for unspecific binding. These experiments showed a significant reduction of the frequency of W2A:W2A as well as WT:W2A interactions compared to WT:WT (Figure 1C). The frequency of the remaining W2A bindings was comparable to those of human Fc interactions, which indicates mostly non-specific interactions. Homotypic WT interactions display a clear peak, while the remaining W2A interaction forces were more spread, again pointing to the non-specificity of these interactions (Figure 1D). The lifetime of DSG2 interactions under zero force was determined by fitting the binding forces detected at different loading rates using Bell’s equation^21^ as done for desmosomal and classical cadherins before, ^22–24^ yielding 1.088 s with R = 0.993. In contrast, no sufficient fit (R = 0.509) was possible for W2A interactions (Figure 1E). Together, these data outline that the tryptophane swap is the major interaction mechanism of DSG2.

To determine the effect of the W2A mutation on cohesion on a cellular level, DSG2-WT and W2A constructs were stably expressed in the epithelial cell line CaCo2 carrying either a WT or DSG2-deficient background [CaCo2(WT) or CaCo2(DSG2 KO), respectively.^25^] Cell-cell dissociation assays were performed, in which a confluent cell monolayer is detached by enzymatic digestion and exposed to defined mechanical stress.^26^ Expression of DSG2-W2A in CaCo2(WT) cells significantly reduced intercellular adhesion as indicated by an increased number of fragments (Figure 1F-H). Importantly, while expressing DSG2-WT in the CaCo2(DSG2 KO) line significantly rescued the impaired intercellular adhesion in response to DSG2 loss, the expression of DSG2-W2A had no effect on fragment numbers. This demonstrates an essential role of the DSG2 tryptophane swap for cell-cell adhesion and points to a dominant negative effect of the DSG2-W2A mutant.

### DSG2-W2A mutant mice resemble the phenotype of ACM

As these data demonstrate the tryptophane swap to be the central adhesive mechanism, we next generated a CRISPR/Cas9 based knock-in mouse model for DSG2-W2A to investigate the consequences of impaired DSG2 adhesion *in vivo* (Figure 2A). Successful generation of founder mutant mice was confirmed by sequencing (Supplementary Figure 1A) and an AluI restriction-based PCR genotyping protocol (Supplementary Figure 1B and method section).

**Figure 2.**
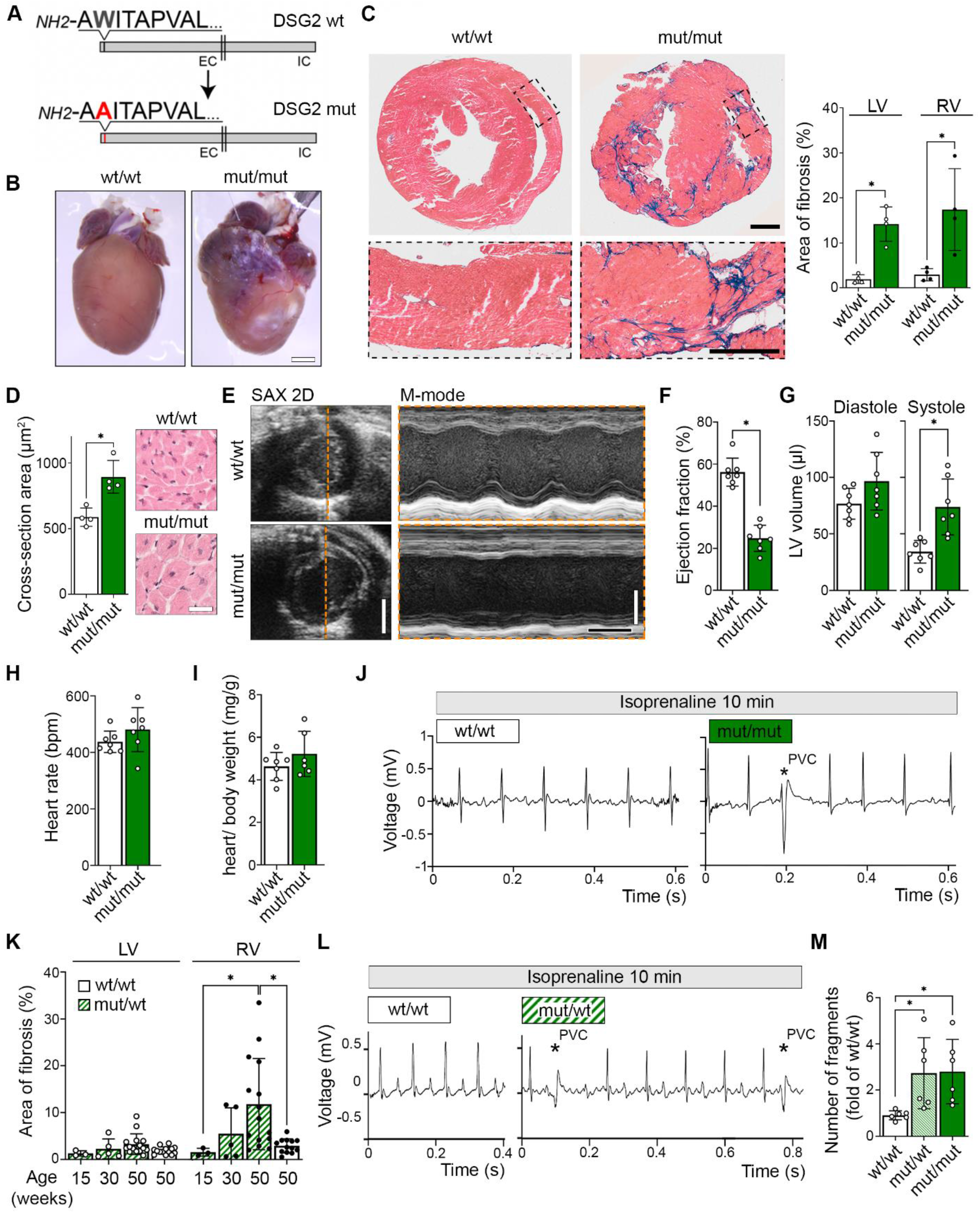
DSG2-W2A mutant mice develop Arrhythmogenic Cardiomyopathy. (**A**) Schematic of the DSG2-W2A mutation within the mature DSG2 protein. The extracellular (EC) and intracellular (IC) part of the molecule is indicated. (**B**) Macroscopic and (**C**) microscopic cardiac phenotype of DSG2-W2A mut/mut mice at the age of 15 weeks. **C** shows a Masson’s trichrome staining with corresponding analysis of the area of fibrosis (blue) in the right (RV) and left ventricle (LV). Black rectangles depict area of zoomed insert. Scale bars: **B** 2 mm; **C** overview: 1 mm; **C** insert: 0.5 mm.*P< 0.05, two-way ANOVA, Sidak’s post hoc test. (**D**) Cardiac hypertrophy was analysed as mean cross-sectional area of cardiomyocytes in Haematoxylin/Eosin stained sections. Scale bar: 30 µm. *P< 0.05, unpaired Student’s t-test. (**E**) Representative image of echocardiography of 30-weeks old wt/wt and mut/mut hearts in short axis (SAX) 2D B-mode and ventricular tracing along the indicated orange line in M-mode. Scale bars: white 2 mm; black 100 ms. (**F, G**) Parameters of left ventricular (LV) function derived from **E**. (**H**) Heart rate of the same mice determined by ECG and (**I**) ratio of heart versus body weight. *P< 0.05, unpaired Student’s t-test. (**J**) Representative ECG of 30-weeks old wt/wt and mut/mut mice after adrenergic stimulation with isoprenaline (2mg/kg body weight i.p.) for 10 min. Asterisk highlights a premature ventricular contraction (PVC). (**K**) Analysis of the area of fibrosis in the right (RV) and left ventricle (LV) in Masson’s trichrome fibrosis staining of DSG2-W2A mut/wt mice at the age of 15, 30 and 50 weeks. *P< 0.05, two-way ANOVA, Sidak’s post hoc test. (**L**) Representative ECG of wt/wt and mut/wt mice at the age of 50 weeks after adrenergic stimulation with isoprenaline (2mg/kg body weight i.p.) for 10 min. Asterisks highlight the PVCs. (**M**) Dissociation assays in immortalized keratinocytes isolated from neonatal murine skin of the respective genotype. *P< 0.05, one-way ANOVA, Dunnett’s post hoc test.

Mating of heterozygous mice revealed a reduction of mut/mut animals’ offspring to 2.8 % compared to the expected Mendelian ratio of 25 %. Dissection of embryos between developmental day E9 and E20 showed loss of the mut/mut animals between E12 and E14 (Supplementary Figure 2A). Macroscopically, mut/mut embryos at day E12.5 appeared pale with accumulation of blood in the cardiac area, while the heart was still beating (Supplementary Figure 2B). On the microscopic level, cells suggestive for blood precursor cells were detectable in the pericardial space of these mice (Supplementary Figure 2C). This finding suggests a rupture of the cardiac wall leading to pericardial bleeding and loss of the animal during development. From the mut/mut mice surviving until birth, an increased percentage of 9.4 % died over an evaluation period of 1 year compared to 0.4 % for mut/wt and 0.5 % for wt/wt animals.

Analysis of adult mut/mut mice at the age of 15 weeks revealed a drastic cardiac phenotype with ventricular deformation, fibrotic and calcified areas (Figure 2B). Large areas of fibrosis were detectable in histological sections of both ventricles and accompanied by hypertrophy of cardiomyocytes in mut/mut hearts (Figure 2C, D). Echocardiography revealed a strong impairment of systolic function with reduced ejection fraction and increased systolic volume, whereas heart rate and weight were not different (Figure 2E–I, Supplementary Figure 3A–D). Baseline ECG during echocardiography displayed no arrhythmias for both genotypes, however, simulation of physical exercise by administration of the adrenergic stimulant isoprenaline for 10 min induced premature ventricular contractions (PVCs) in 28.6 % (2 out of 7) of the mut/mut animals, while wt/wt mice developed no arrhythmias (Figure 2J). Together, DSG2-W2A mut/mut animals displayed left ventricular systolic dysfunction, biventricular fibrosis and occasional PVCs under pharmacologically induced stress (isoprenaline). In comparison to the Padua criteria for the diagnosis of ACM in patients,^27^ DSG2-W2A mut/mut animals resemble the phenotype of biventricular ACM fulfilling the equivalent of a minimum of two major and one minor criteria. However, in ACM patients the majority of mutations is hemizygous^17^. In contrast to homozygous mutant animals, mut/wt DSG2-W2A animals showed no fibrosis at the age of 15 weeks. However, increasing fibrosis mainly of the right ventricle was detectable in mut/wt mice until the age of 50 weeks (Figure 2K, Supplementary Figure 3E). Interestingly, the penetrance of this effect appeared to be heterogeneous, as in 26.7 % of the 50-weeks old mut/wt animals percentage of fibrotic areas was similar to wild type mice. Simulating physical stress by isoprenaline, 50 % (5 out of 10) of the heterozygous animals at the age of 50 weeks developed PVCs (Figure 2L). In summary, DSG2-W2A mut/wt animals show progressive fibrosis of the right ventricle and ventricular arrhythmia. Thus, according to the Padua criteria,^27^ the heterozygous genotype mimics the characteristics of a right-dominant ventricular ACM by fulfilling the equivalent of a minimum of two major criteria. To confirm loss of cell-cell adhesion by the DSG2-W2A mutation in this mouse model, keratinocytes were isolated from the skin of neonatal animals and subjected to dissociation assays. These experiments showed a significant increase in monolayer fragmentation in cells derived from mut/wt and mut/mut mice compared to wt/wt (Figure 2M). Together, these data demonstrate that loss of desmosomal adhesion is sufficient to induce ACM and that the DSG2-W2A model can resemble two different phenotypes of the disease.

### Junctional proteins are preserved at the ICD of DSG2-W2A mutants

We next applied this mouse model to investigate mechanisms leading to the ACM phenotype. RNA sequencing was performed from ventricles of mut/mut and wild type hearts both before onset of fibrosis (age of 5 days), and when the biventricular fibrosis and ACM phenotype was well established (after 9 weeks). Principal component analysis (PCA) showed sufficient clustering of the respective sample groups (Supplementary Figure 4A). To validate that the DSG2-W2A mouse model is resembling gene expression similar to ACM, sequencing data from 9-week old mice with ACM phenotype were compared to the top differentially expressed genes derived from published transcriptomic data sets of ACM patient hearts (GEO data base, GSE107157/GSE107480 and GSE29819^28^). Murine samples clustered according to their genotype and mut/mut hearts resembled the gene expression pattern of patients (Supplementary Figure 4B). Together, this proves the reliability and comparability of the sequencing data from DSG2-W2A mice to patients.

Interestingly, RNA-Seq of DSG2-W2A mutant hearts demonstrated largely unchanged expression levels of individual junctional adhesion molecules (Supplementary Figure 5A). As posttranslational processes including molecule turn-over and correct subcellular distribution are important regulators of mechanical junctions,^29, 30^ we investigated protein expression and localization of desmosomal and adherens junction components in adult murine hearts. These experiments revealed a reduction of DSG2 protein levels in mutant hearts with the remaining molecules properly localizing to the ICD, while no other alterations of junctional molecules could be detected (Supplementary Figure 5B, C). In addition, the mRNA expression levels of gap and tight junction components, which are also part of the ICD, showed no consistent changes (Supplementary Figure 5D, E). These data indicate that the majority of junctional components are well preserved at the ICDs of DSG2-W2A mutant hearts.

### Deregulation of Integrin-β6 in ACM patient and DSG2-W2A mouse samples

To identify common genes deregulated during ACM pathogenesis, we compared the significantly altered genes from patient data sets with the results from mutant mice. As we were mainly interested in mechanisms inducing ACM, we included data from 5-days old mice to identify changes already present before onset of secondary effects due to fibrosis. Interestingly, integrin-β6 (*Itgb6*) was the only gene consistently deregulated in all data sets (Figure 3A). Its common down-regulation in ACM patients and mutant mice on RNA levels was further confirmed for heterozygous DSG2-W2A mutants compared to wild type by quantitative RT-PCR (Figure 3B). Surprisingly, however, protein levels were significantly increased in adult mut/mut hearts (Figure 3C). This was further supported by immunostaining, which showed enhanced intensity of ITGB6 at the membrane of mutant cardiomyocytes (Figure 3D). This suggests that the reduction on mRNA level is a compensatory mechanism for the increased protein levels. Moreover, ITGB6 staining was not only detectable at the ICD in mutant animals but intensity also increased at the lateral membrane of the cell. This effect was even more pronounced adjacent to fibrotic areas. In addition to mouse samples, cardiac tissue of an ACM patient bearing a DSP-E952X mutation, who died from this disease at the age of 14, was analysed with respect to ITGB6. Importantly, compared to healthy controls, patient cardiomyocytes also displayed increased ITGB6 staining at the ICD and lateral membrane (Figure 3E). In line with a stop-codon leading to a truncated DSP, the protein was not detectable in patient tissue. Together, these data demonstrate a deregulation of ITGB6 in ACM patients and DSG2-W2A hearts with down-regulation on RNA but up-regulation on protein level together with alteration in molecule distribution.

**Figure 3.**
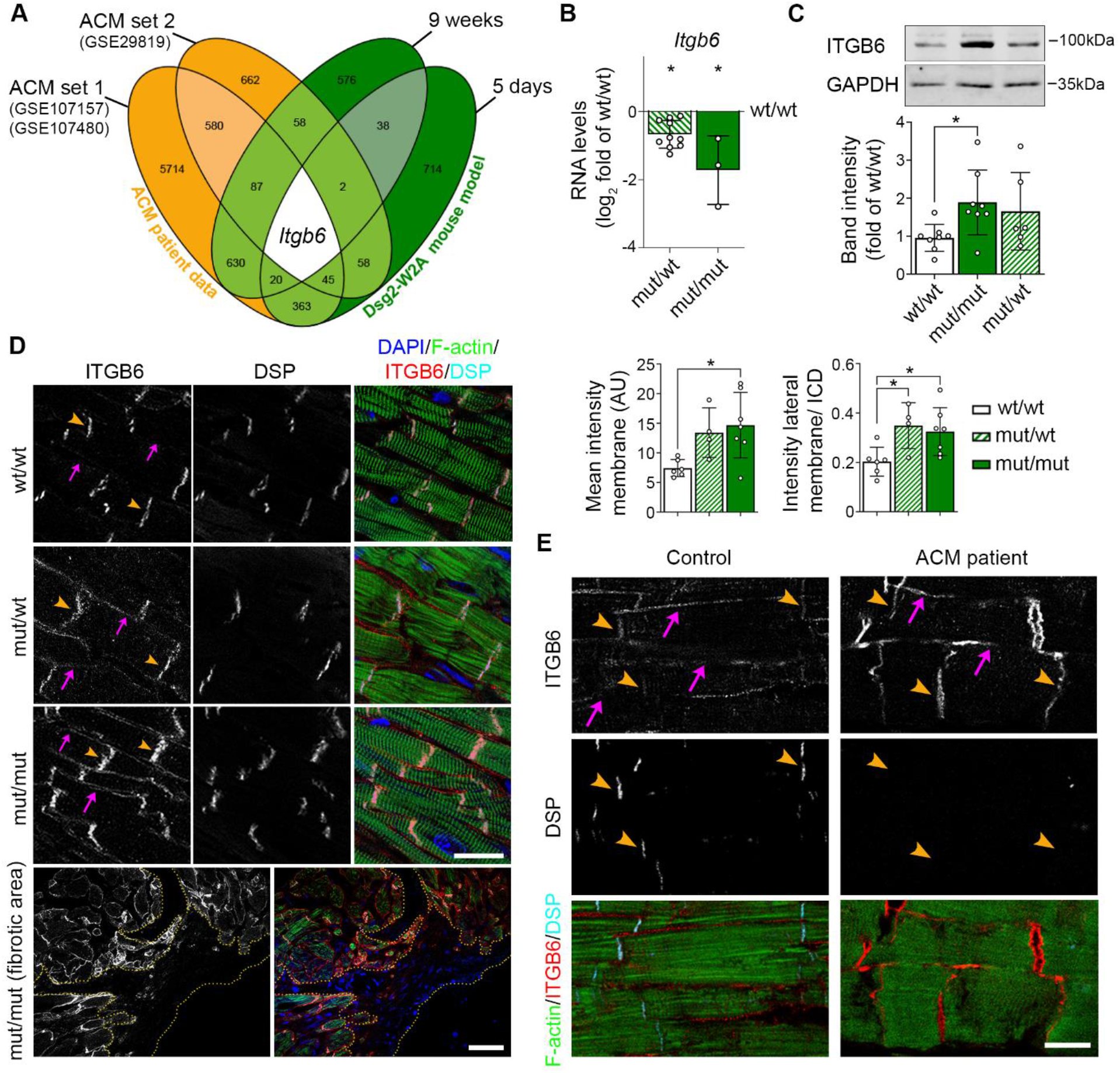
Integrin-β6 is deregulated in DSG2-W2A mutants. (**A**) Venn diagram of significantly altered genes from indicated ACM patient data sets (ACM vs. healthy control) and DSG2-W2A mice at the age of 5 days and 9 weeks (mut/mut vs. wt/wt) highlighting integrin-β6 (*Itgb6*) as only overlapping gene with same direction of expression in all data sets. Numbers indicate the amount of overlapping genes for the respective overlays. (**B**) RNA expression of *Itgb6* analysed via q-RT-PCR in adult DSG2-W2A mouse hearts. *P< 0.05, unpaired Student’s t-test vs. wt/wt. (**C**) Representative Western blot and respective analysis of band intensity of ITGB6 in DSG2-W2A hearts. GAPDH served as loading control. *P< 0.05, one-way ANOVA, Dunnett’s post hoc test. (**D**) Immunostaining of ITGB6 (red in overlay) in DSG2-W2A hearts with corresponding analysis of staining intensity at membrane region in total and ratio of staining intensity at the lateral membrane (pink arrows) vs. ICD area (orange arrow heads). Desmoplakin (DSP, cyan) marks ICDs, DAPI (blue) nuclei and F-actin (green) the sarcomere system. Lower row shows an overview image of a fibrotic area in mut/mut hearts. Dotted orange line outlines the fibrotic area. Scale bars: upper rows: 20 µm, lower row: 50 µm. *P< 0.05, one-way ANOVA, Dunnett’s post hoc test. (**E**) Representative immunostainings of ITGB6 (red) and DSP (cyan) in an ACM patient (DSP-E952X) and a healthy control sample. Lateral membrane (pink arrows) and ICDs (orange arrow heads) are highlighted. F-actin (green) stains the sarcomere system. For the ACM patient, 4 different tissue samples were analysed and compared to 2 tissue samples from 2 healthy controls. Scale bar: 20 µm.

### Spatial organisation and interaction of integrin-β6 and DSG2 at the ICD

As these data suggest modulation of ITGB6 on posttranslational level, we investigated the spatial distribution of this protein in more detail using structured illuminated microscopy (SIM). Image stacks of ICDs, where the majority of the ITGB6 was present, demonstrated clustering in distinct regions throughout the junctional interface of wild type hearts (Figure 4A). In contrast, mutant cardiomyocytes displayed a more homogeneous distribution of ITGB6 signals at the ICD without clear grouping, which was objectified as increased mean distance to the nearest neighbours. Numbers or volume of individual ITGB6 signals was not changed under these conditions. As this morphological alteration suggests changes in the junctional ultrastructure, we performed transmission electron microscopy (TEM). These experiments revealed disturbed ICDs with widened intercellular space and occasionally completely ruptured junctions in mut/mut hearts (Figure 4B). Further, SIM analysis demonstrated reduced number and size of DSG2 signals at W2A mutant ICDs (Figure 4C). Together, this suggests that reduced DSG2 levels lead to disturbance of the ICD structure and ITGB6 localization in W2A cardiomyocytes. Next, we assessed the possibility that DSG2 regulates ITGB6 localization and clustering by direct protein-protein interaction. DSG2 was pulled down together with ITGB6 in co-immunoprecipitation experiments in wild type hearts extracts (Figure 4D). In mutant heart lysates, interaction of ITGB6 with DSG2 was clearly reduced and SIM experiments confirmed a diminished spatial proximity of ITGB6 signals to DSG2 in the ICD (Figure 4E). Nevertheless, the remaining DSG2 molecules localized correctly to ITGB6. In summary, these data point to reduced structural interactions between ITGB6 and DSG2 as consequence of the diminished DSG2 levels, which prevents the clustering of ITGB6 at the ICD.

**Figure 4.**
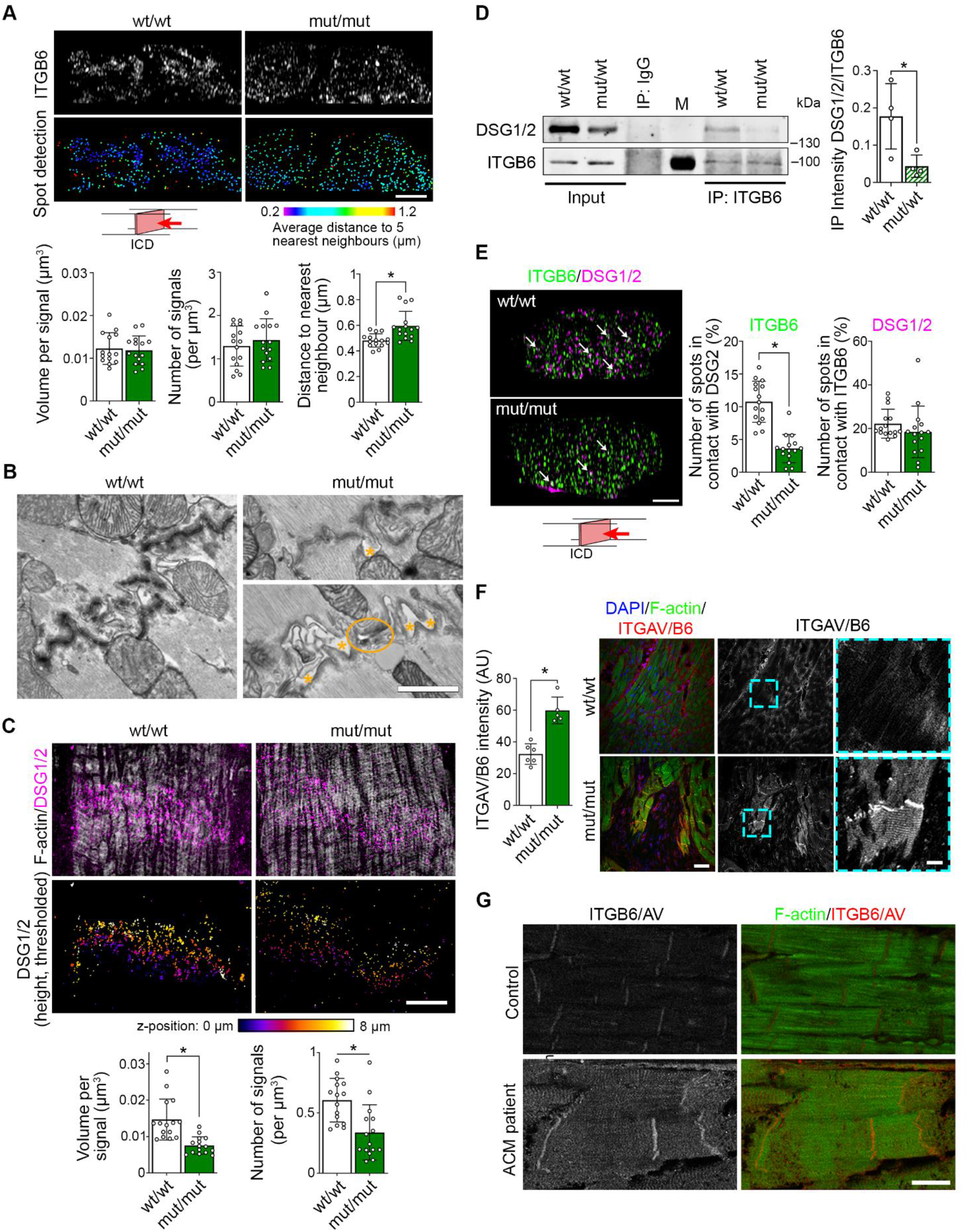
Altered spatial distribution of ITGB6 at the ICD of DSG2-W2A mutant mice. (**A**) Representative ICD reconstruction from z-stacks of ITGB6 immunostaining acquired by structured illumination microscopy (SIM). Lower row shows signal spot detection with indication of the average distance to the 5 nearest neighbour signals. Pictogram displays the angle of view. Related analysis of ITGB6 signal volume, number and average distance to the five nearest neighbours is shown below. Scale bar: 2 µm. *P< 0.05, unpaired Student’s t-test. Each dot represents the mean value of one ICD from in total 3 mice per genotype. (**B**) Representative images of ICDs acquired by transmission electron microscopy (TEM) with 3 mice per genotype. Orange asterisks mark intercellular widening, orange circle marks a ruptured junction. Scale bar: 1 µm. (**C**) Representative images of DSG2 (magenta) and filamentary actin (f-actin, white) stacks acquired by SIM and presented as z-stack maximum intensity projection. Lower row shows color-coded height projection of DSG2 signals in z-stack after signal thresholding as performed for analysis. Related analysis of DSG2 signal volume and number is shown below. Scale bar: 5 µm. *P< 0.05, unpaired Student’s t-test. Each dot represents the mean value of one ICD from in total 3 mice per genotype. (**D**) Representative immunoblot from immunoprecipitation (IP) of ITGB6 and DSG2 co-immunoprecipitation with analysis on the right. Intensity of co-immunoprecipitated DSG2 was normalized to the amount of pulled-down ITGB6. IP of IgG of the same species as anti-ITGB6 served as control for unspecific binding, M marks the lane with height marker. *P< 0.05, unpaired Student’s t-test. (**E**) Representative ICD reconstruction from z-stacks of ITGB6 (green) and DSG1/2 (magenta) immunostaining acquired by SIM. Analysis of the fraction of ITGB6 signals in close contact (< 200 nm) to a DSG2 signal and vice versa is shown on the right. Arrows highlight examples of colocalizing signals (white). Scale bar: 2 µm. *P< 0.05, unpaired Student’s t-test. Each dot represents the mean value of one ICD from in total 3 mice per genotype. (**F**) Immunostaining of ITGAV/B6 heterodimer in DSG2-W2A mutant hearts with respective analysis of staining intensity. Cyan rectangle marks zoomed area on the right. Scale bars: overview 50 µm; insert 10 µm. *P< 0.05, unpaired Student’s t-test. (**G**) Representative immunostaining images of ITGAV/B6 heterodimer staining (red) in a ACM patient (DSP-E952X) and healthy control sample. F-actin (green) stains the sarcomere system. For the ACM patient, 4 different tissue samples were analysed and compared to 2 tissue samples from 2 healthy controls. Scale bar: 20 µm.

### Increased interaction of integrin-β6 and integrin-αV

ITGB6 needs to hetero-dimerize with integrin-αV (ITGAV) in order to be activated and bind to the extracellular matrix.^31–33^ Thus, we analyzed the expression of the counterpart ITGAV, which was significantly upregulated on protein level and increased at the ICD and lateral membrane of mutant hearts (Supplementary Figure 6A, B). In contrast, the localization and intensity of integrin-β1 (ITGB1), as classical representative of the integrin group, at the lateral membrane was not altered (Supplementary Figure 6C). Importantly, staining with an antibody specifically recognizing the heterodimer of ITGAV/B6 revealed increased amount of dimers at the ICD and lateral membrane in mutant mice (Figure 4F). A comparable effect with enhancement of ITGAV/B6 staining intensity was further detectable in the ACM patient sample (Figure 4G). These data suggest a sequence by which impaired junctional integrity due to loss of DSG2 binding function leads to an increase and delocalization of ITGB6, which can then dimerize to larger extents with ITGAV to switch to its active state.

### Upregulation of TGF-β signaling in DSG2-W2A mutants

Importantly, the ITGAV/B6 dimer has the ability to activate the pro-fibrotic cytokine TGF-β by binding and removal of the latency associate peptide (LAP).^31, 32^ Gene set enrichment analyses revealed an upregulation of genes associated with TGF-β signaling in both 9-weeks old DSG2-W2A mutants as well as ACM patients (Figure 5A). Moreover, direct targets of receptor-regulated SMADs (R-SMADs) involved in TGF-β signaling according to the TRRUST data base^34^ were upregulated in DSG2-W2A mutant hearts (Figure 5B). Accordingly, increased amounts of nuclear SMAD2/3 phosphorylated at the activation sites S465/S467 or S423/S425, respectively, were detected in these hearts (Figure 5C). Interestingly, increased levels of pSMAD2/3 were not only found in fibroblasts but also in cardiomyocytes mainly adjacent to fibrotic areas.

**Figure 5.**
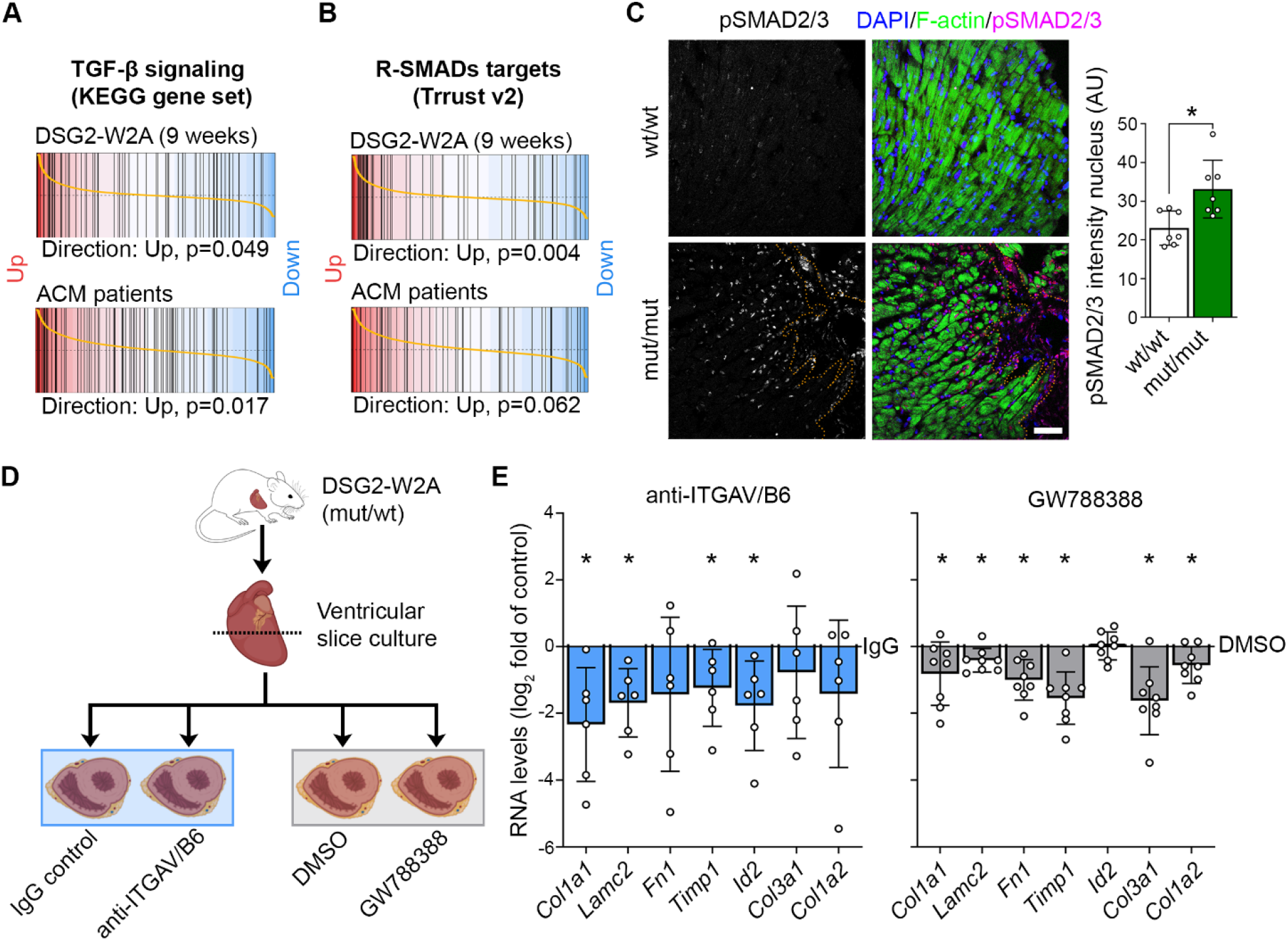
Elevated TGF-β signaling in DSG2-W2A hearts as result of ITGAV/B6 activity. Barcode blots of gene set enrichment analysis of (**A**) the KEGG_TGF_BETA_SIGNALING_PATHWAY data set (systematic name: M2642,^56^ or (**B**) genes directly regulated by receptor-regulated SMADs (R-SMADs, includes SMAD1/2/3/5/9) as published in the TRRUST data base^34^ in 9-weeks old DSG2-W2A (mut/mut vs wt/wt) or ACM patient data set 1 (ACM vs. healthy control, GEO: GSE107157/GSE107480). (**C**) Immunostainings of phosphorylated SMAD2/3 (magenta, S465/S467 or S423/S425, respectively) in sections of DSG2-W2A hearts and related analysis of nuclear staining intensity. Nuclei are stained with DAPI (blue), cardiomyocytes are marked with f-actin (green). Dotted orange line marks edge of fibrotic area. Scale bar: 50 µm. *P< 0.05, unpaired Student’s t-test. (**D**) Schematic of experimental set-up for ITGAV/B6 blocking experiments in cardiac slice culture with related results in **E**. Icons are derived from BioRender. (**E**) q-RT-PCR analysis of expression of genes downstream of TGF-β signaling in cardiac slices cultures treated with inhibiting anti-ITGAV/B6 (1:15) or 10 µmol/l GW788388, an inhibitor of TGF-β receptor I, for 24 hours. *P< 0.05, paired Student’s t-test vs. indicated control condition.

### Downregulation of TGF-β target genes by ITGAV/B6 inhibition

These data indicate upregulation of the pro-fibrotic TGF-β pathway and its downstream targets, which include extracellular matrix proteins such as collagens, laminin or fibronectin. Because the ITGAV/B6 dimerization is increased in DSG2-W2A and known to activate TGF-β molecules, we investigated whether inhibition of ITGAV/B6 was efficient to reduce expression of fibrosis markers in DSG2-W2A mice. For these experiments, cardiac slice cultures were generated from the ventricles of heterozygous mice at the age of 40-50 weeks and treated for 24 hours with anti-ITGAV/B6 (clone 10D5), which was shown to neutralize the function of the dimer *in vitro* as well as *ex vivo*.^35, 36^ In comparison, slices were treated with the TGF-β receptor I (ALK5) inhibitor GW788388^37^ to directly block TGF-β signaling. Heterozygous mice were chosen, as they resemble the most common phenotype in ACM patients (Figure 5D). As revealed by quantitative RT-PCR, significant downregulation in the expression of the pro-fibrotic molecules α1 type I collagen (*Col1a1*), laminin subunit-γ2 (*Lamc2*), metalloproteinase inhibitor-1 (*Timp1*), and ID-2 (*Id2*) was induced in the right ventricle of mutant mice in response to inhibition of ITGAV/B6 (Figure 5E). Other extracellular matrix proteins such as fibronectin (*Fn1*) or collagens (*Col3a1, Col1a2*) showed the same tendency. Importantly, all of these fibrotic markers were significantly upregulated in adult mutant hearts as detected by RNA-Seq (Figure 5A, B). Further, direct inhibition of TGF-β led to a similar downregulation of these markers as the blocking antibody. In conclusion, these experiments link the upregulation of the ITGAV/B6 dimer in response to abrogated DSG2 binding to a pro-fibrotic expression pattern induced by TGF-β. Moreover, they outline ITGAV/B6 as potential treatment target to inhibit cardiac fibrosis.

## Discussion

In this study, we generated a knock-in mouse model with defective binding function of the adhesion molecule DSG2, which demonstrated that defective desmosomal adhesion is sufficient to induce an ACM phenotype comparable to human disease. By RNA sequencing of DSG2-W2A mouse hearts at different time points of disease progression and comparison of the results with ACM patient data identified integrin-β6 (ITGB6) as commonly altered molecule. Interestingly, subsequent super-resolution immunostaining and immunoprecipitation revealed reduced interaction of ITGB6 with DSG2 in mutant hearts. Our data further suggest an increased dimerization of ITGB6 with its counterpart ITGAV and activation of TGF-β signaling with expression of pro-fibrotic proteins such as collagen-1 and laminin as consequence. Importantly, antibody-mediated inhibition of ITGAV/B6 was sufficient to reduce expression of these molecules in cardiac tissue. In summary, our data demonstrate a cascade of defective desmosomal adhesion and disrupted ICD structure to activation of ITGAV/B6 with pro-fibrotic TGF-β signaling as important underlying mechanism leading to an ACM phenotype (Figure 6). Finally, we show that this pathway can be targeted by drug treatment, which highlights the relevance for development of future therapeutic options in patients.

**Figure 6.**
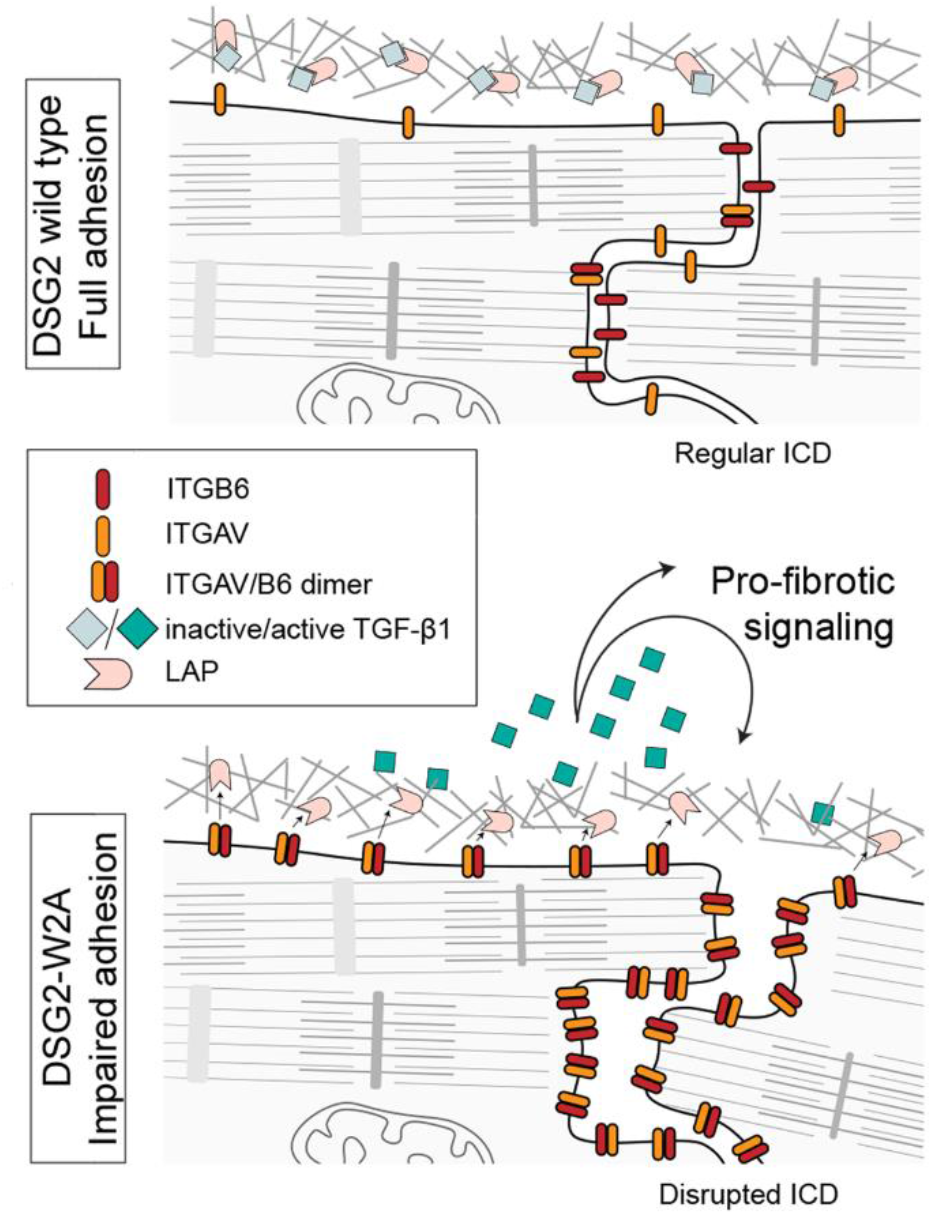
Schematic conclusion of data. DSG2-W2A mutation with loss of desmosomal adhesion leads to impaired ICD structure with deregulation of ITGB6 and enhanced heterodimerization with ITGAV. The dimer efficiently binds to the extracellular matrix and activates TGF-β by removal of the latency-associated peptide (LAP). Active TGF-β can then induce pro-fibrotic downstream signaling via SMAD molecules. In our experiments, this cascade was blocked by anti-ITGAV/B6 or TGF-β receptor I inhibitor GW788388.

### Defective desmosomal adhesion by DSG2-W2A mutation is inducing ACM

Due to mutations mainly affecting desmosomal genes and evidence of disrupted ICDs, the hypothesis of dysfunctional desmosomes with loss of cell-cell adhesion as central pathological step was soon adopted in the field.^2, 17^ However, experimental data on this topic are contradictory.^13–16^ Applying force spectroscopy, this study provides evidence that the tryptophane swap is the major binding mechanism of DSG2, as predicted by crystal structure.^19^ Further, cell-cell dissociation experiments showed that this interaction is essential for effective intercellular adhesion. By generating a DSG2-W2A knock-in mouse model, we here show that disrupted desmosomal adhesion is sufficient to induce an ACM phenotype fulfilling the Padua criteria^27^ used for diagnosis in patients including ventricular fibrosis, arrhythmia, and impaired ventricular function. In line with this, three mutations are described in ACM patients directly affecting the DSG2 binding mechanism by exchange of the tryptophane with a serine,^38^ leucine,^39^ or arginine (ClinVar data bank, variation ID 420241). This indicates a disruption of the tryptophane swap independent from the substituting amino acid and supports defective desmosomal adhesion as important factor in ACM.

### DSG2-W2A mice as model to mimic ACM

As outlined above, by precisely targeting the central binding motif of DSG2, our data provide strong support for the hypothesis of impaired cardiomyocyte cohesion as crucial step towards ACM. In line with this, another model, in which large parts of the two outermost DSG2 extracellular domains were deleted (but leaving the W2A section intact) showed a similar phenotype.^40^ Thus, the DSG2-W2A mouse model reproduces a common pathogenic mechanism (i.e. reduced intercellular adhesion of cardiomyocytes) occurring independently from the underlying mutation. As ACM can be induced by mutations in different desmosomal and non-desmosomal molecules, this model can serve as an excellent tool to study joint mechanisms and avoid effects specific to a distinct mutation or protein. Further, the model is based on a knock-in with exchange of a single amino acid under the endogenous promotor. This reduces the possibility of secondary unwanted effects due to complete protein absence or overexpression under an exogenous promotor as performed in other ACM models. Similar to patients, the mutation is present in all cardiac cell types and not limited to cardiomyocytes.^2, 18^ Comparison of transcriptomic data demonstrated comparable expression patterns in the DSG2-W2A mouse model and ACM patients bearing mutations in a variety of genes.^28^ Furthermore, effects on ITGAV/B6 found in DSG2-W2A mice were also detectable in ACM patient sample with a DSP mutation. This highlights the DSG2-W2A mouse model as a relevant tool to mimic this disease independent from the underlying mutation.

### DSG2-W2A affects ITGAV/B6

Investigating the mechanism downstream of impaired adhesion, we identified ITGB6 as molecule commonly deregulated in ACM patients and DSG2-W2A mice. Especially the inclusion of DSG2-W2A mutant hearts before and after onset of disease in the comparison with ACM patient data points to a basic role of ITGB6 in disease development before establishment of secondary effects such as fibrosis. Even though transcriptomic data and q-RT-PCR revealed a consistent downregulation on RNA level, the protein was significantly upregulated in Western blot and immunostaining analysis. This can be interpreted as compensatory attempt to balance the amount of a protein, which might be more stable under mutant conditions. The increased levels of ITGB6 protein were accompanied by a more uniform distribution within the ICD and additional signals present at the lateral membrane. These changes were paralleled by a specific reduction of DSG2 at the ICD with less and smaller signals, a reduced interaction and spatial proximity of DSG2 and ITGB6 as detected by immunoprecipitation and super-resolution imaging and general impaired integrity of the ICD as demonstrated by electron microscopy. These results are consistent with a model in which reduced stability of DSG2 at the membrane due to impaired binding is hampering sequestration of ITGB6 at the ICD. This reduced sequestration might be important for the availability of ITGB6 for dimerization with ITGAV and corresponding downstream signaling. So far, only limited data are available on the role of integrins in ACM. A recent study showed downregulation of integrin-β1D in ACM, which is suggested to cause ventricular arrhythmia involving ryanodine receptor 2 phosphorylation and aberrant calcium handling.^41^ Further, knock-down of PKP2 in HL-1 cardiomyocytes was described to deregulate focal adhesions including integrin-α1.^42^ Thus, our study uncovers a role of ITGAV/B6 in ACM pathogenesis and demonstrates a novel regulation of ITGAV/B6 distribution and activity by DSG2 binding.

### Activation of TGF-β signaling by ITGAV/B6 as mechanism in ACM

In this study, deregulation of ITGB6 correlates with elevated levels of ITGAV and increased hetero-dimerization of both molecules. Because ITGAV is described as exclusive integrin binding partner of ITGB6, while ITGAV can interact with other β-integrins, ITGB6 is assumed to control ITGAV/B6 levels.^33, 43^ ITGAV/B6 binds to the RGD peptide motif present in extracellular matrix proteins such as fibronectin or the latency-associated peptide (LAP), which is part of pro-*TGF-*β. By mechanical force exerted through ITGAV/B6, TGF-β is detached from LAP and can then induce signaling via TGF-β receptors and the SMAD cascade.^32, 44^ With this mechanism, the *ITGAV/B6* hetero-dimer is described as one of the major activators of latent TGF-β1 and TGF-β3 in vivo.^32, 44^ Further, regulation of TGF-β receptor II by the cytoplasmic tail of ITGAV/B6 was described.^45^

The functional connection of ITGAV/B6 and TGF-β is of special importance for cardiomyopathies, as TGF-β signaling is known as general driver of cardiac fibrosis ^46, 47^ and more specifically was implicated in ACM.^2, 48, 49^ Even though the implication of ITGAV/B6 for pro-fibrotic signaling and its impact as therapeutic target are known from other diseases such as liver, kidney, or pulmonary fibrosis, to our knowledge no data are available on the role of these molecules and their regulation of TGF-β signaling in cardiac fibrosis.^50^ This highlights the relevance of this study, revealing a deregulation of ITGB6 under different ACM conditions with subsequent upregulation of a pro-fibrotic expression pattern. Uncovering this pathway is of high interest, as it offers the possibility to target TGF-β with reduced risk of severe side effects occurring in response to direct TGF-β inhibition. ^51^ Inhibition of ITGAV/B6 by small molecules such as EMD527040 and GSK3008348 or neutralizing antibodies was shown to be protective in murine models of lung, liver and biliary fibrosis.^52–55^ Accordingly, we demonstrate an effective reduction of pro-fibrotic gene expression under ACM conditions in response to treatment with an ITGAV/B6 inhibiting antibody.

In conclusion, we established a new ACM mouse model and uncovered a novel pathway of fibrosis induction. Furthermore, we identified an approach to target this mechanism with future implication as potential therapeutic option in patients.

## Methods

### Human heart samples

Heart samples of ACM patient and healthy controls were derived from forensic autopsy. This study was conducted according to the tenets of the Declaration of Helsinki and approved by the local ethic committees, licence number 494-16 (ethic committee LMU Munich, Germany) and 152/15 (ethic committee Goethe University, Frankfurt am Main, Germany). Samples were fixed and embedded in paraffin according to standard procedures.

### Mouse experiments and DSG2-W2A mouse model

All mouse experiments were carried out according to the protocol approved by the Cantonal Veterinary Office of Basel-Stadt (License number 2973_32878 and 3070_32419). All mice were housed under specific pathogen-free conditions according to institutional guidelines. Animals of both sexes were applied without bias.

The Dsg2-W>A allele was obtained by Cas9/CRISPR embryo electroporation. The Cas9/CRISPR target sequence tggttcgtcaaaagagggcc(tgg) (PAM sequence in brackets is also the TGG-Trp codon) spanning the mutation site was selected with the help of CRISPOR software (http://crispor.tefor.net/)56. ssDNA oligonucleotide 5’gtgataactcaaggtaattgtattaacaggtcttcagcccaagaaatgaaggcaaaccgttccctaagcacactcac ttggttcgtcaaaagagggca**<u>gct</u>**atcactgcccctgtggctctgcgggagggcgaagacctgtccagaaagaacc cgattgccaaggtagcagctacagaagaatgtggcgagggtgttggc3’ (GCT – Ala codon in bold underlined) was designed to insert the W>A mutation into the Cas9-generated DSB by homologous recombination and at the same time mutate the TGG PAM sequence to GCT. C57BL/6J female mice underwent ovulation induction by i.p. injection of 5 IU equine chorionic gonadotrophin (PMSG; Folligon, InterVet, Vienna, Austria), followed by i.p. injection of 5 IU human chorionic gonadotropin (Pregnyl, Essex Chemie, Lucerne, Switzerland) 48 hours later. For the recovery of embryos, C57BL/6J females were mated with males of the same strain immediately after the administration of human chorionic gonadotropin. Embryos were collected from oviducts 24 hours after the human chorionic gonadotropin injection, and were then freed from any remaining cumulus cells by a 1–2 min treatment of 0.1 % hyaluronidase (Sigma-Aldrich, St. Louis, MO, ISA) dissolved in M2 medium (Sigma-Aldrich). Prior to electroporation, the zona pellucida was partially removed by brief treatment with acid Tyrode’s solution and the embryos were washed and briefly cultured in M16 medium (Sigma-Aldrich) at 37 °C and 5 % CO_2_. Electroporation with a mixture of ssDNA oligonucleotide targeting template, 16 µmol/l cr:trcrRNA hybrid targeting Dsg2 and 16 µmol/l Cas9 protein (all reagents from IDT, Coralville, IA, USA) was carried out using 1 mm gap electroporation cuvette and the ECM830 electroporator (BTX Harvard Apparatus, Holliston, MA, USA). Two square 3 ms pulses of 30 V with 100 ms interval were applied as previously described.^57^ Surviving embryos were washed with M16 medium and transferred immediately into the oviducts of 8–16-weeks-old pseudopregnant Crl:CD1(ICR) females that had been mated with sterile genetically vasectomized males ^58^ the day before embryo transfer (0.5 dpc). Pregnant females were allowed to deliver and raise their pups until weaning age. In total 150 embryos were electroporated and 147 surviving embryos were transferred into 7 foster mothers. All foster mothers produced live litters with a total of 20 viable F0 pups. One F0 pup carried the desired mutation as confirmed by sequencing. This founder animal was bred to C57BL/6J partner. The mut/wt offspring from this mating was bred to C57BL/6J partners for 2 generations to establish the Dsg2-W2A mouse line.

For genotyping of the DSG2-W2A line, DNA was extracted from biopsies in 25 mmol/l NaOH and 0.2 mmol/l EDTA at 98 °C for 1 hour and neutralized with 40 mmol/l Tris pH 5.5. PCR was performed using GoTaq G2 (M7845, Promega, Madison, WI, USA) according to manufacturer’s instructions with the primers Dsg2-W2A for: GAATGTCTCCCCAAAGCTTTGGGTATG and Dsg2-W2A rev: CTGCTACCTTGGCAATCGGGTTC, which span the mutated region. The PCR product was restricted with 66.7 U/ml AluI (R0137, New England Biolabs, Ipswich, MA, USA) in CutSmart buffer (New England Biolabs) overnight at 37 °C. By subsequent electrophoresis in a 3 % (w/v) agarose gel containing Midori Green Advanced (Nippon Genetics, Düren, Germany) for fluorescence DNA visualization, presence of a Dsg2-WT allele was detectable as 197 bp fragment, while Dsg2-W2A mutant allele was cut into a 109 bp and 72 bp fragment (Supplementary Figure 1B).

For heart dissection, mice were euthanized via i.p. pentobarbital overdose according to guidelines of the Cantonal Veterinary Office of Basel-Stadt and the University of Basel. Hearts were removed by lateral thoracotomy and directly immersed in ice-cold HBSS supplemented with 20 mmol/l 2,3-Butanedione monoxime (BDM, Sigma-Aldrich) unless stated otherwise. Morphology of the hearts was analyzed using a binocular stereo microscope (SZX2, Olympus, Shinjuku, Japan) and documented with a SLR camera (EOS 800D, Canon, Tokyo, Japan). Tissue was processed as described in the respective section.

For dissection of embryos, timed matings were performed and pregnant mice euthanized via i.p. pentobarbital overdose after the respective days. Embryos were dissected from the uterus and placed in HBSS. After image acquisition as described above, a tissue sample for genotyping was collected from the tail and embryos were processed as described in the *Histological staining* section.

### Plasmid generation and cloning

For lentiviral overexpression plasmids, DNA for full length Dsg2-WT and full length Dsg2-W2A mutation, respectively, were amplified from cDNA originating from liver tissue of DSG2-WT and DSG2-W2A mice using AscI-msDsg2-N forward and NotI-GT-msDsg2-C reverse primers. Amplicons were ligated into AscI and NotI digested pLENTI-C-mGFP (#PS100071, OriGene, Rockville, MD, USA) according to standard procedures. To produce the proteins used in the AFM experiments, the signal, pro-peptide, and all extracellular domains of Dsg2-WT and Dsg2-W2A, respectively, were amplified from murine cDNA using AfeI-Kozak-msDsg2-N forward and XhoI-msDsg2-C reverse primers. Amplicons were ligated into AfeI and XhoI digested Fc-His-pEGFP-N3 plasmid containing the Fc domain from human IGHG1 (bases 295-990) (a kind gift of Nikola Golenhofen, Institute of Anatomy and Cell Biology, University of Ulm, Ulm, Germany). For the Fc control construct, the signal peptide of murine interleukin 2 was inserted N-terminally of the human Fc by annealing the oligomers NheI-Kozak-IL2 Signal and XhoI-IL2 Signal and ligating them into the NheI and XhoI digested Fc-His-pEGFP-N3 plasmid.

### Primers and oligomers for cloning

AscI-msDsg2-N forward: GTTTGGCGCGCCATGGCGCGGAGCCCGGGT

NotI-GT-msDsg2-C reverse: GTTTGCGGCCGCGTGGAGTAAGAATGTTGCATGGTG

AfeI-Kozak-msDsg2-N forward: GTTTAGCGCTGCCACCATGGCGCGGAGCCCGGGTGA

XhoI-msDsg2-C reverse: GTTTCTCGAGGGCAGGGCCCAACCCGAC

NheI-Kozak-IL2 Signal: CTAGCCACCATGTACAGGATGCAACTCCTGTCTTGCATTGCACTAAGTCTTGCACTTGTCACGAATTCGC

XhoI-IL2 Signal: TCGAGCGAATTCGTGACAAGTGCAAGACTTAGTGCAATGCAAGACAGGAGTTGCATCCTGTACATGGTGG

### Cultivation of CaCo2 cells

The human intestinal cell line CaCo2 with WT and DSG2 KO background were kindly provided by Nicolas Schlegel (Department of General, Visceral, Vascular and Pediatric Surgery, University Hospital Würzburg, Würzburg, Germany) and generated as described.^25^ CaCo2 cells were maintained in Dulbecco’s Modified Eagle Medium (DMEM, D6546, Sigma-Aldrich) supplemented with 10 % foetal bovine serum (S0615, Merck, Darmstadt, Germany), 100 μg/ml penicillin/streptomycin (Applichem, Darmstadt, Germany) and 2 mmol/l L-glutamine (Sigma-Aldrich) at 37 °C, 5 % CO_2_ and full humidity. For experiments, cells were seeded on TC-treated plastic cell culture plates, grown to confluency and differentiated for seven days. All cells were quarterly checked for mycoplasma infections using PCR and were proven negative. CaCo2 cells were routinely authenticated by Short Tandem Repeat profiling.

### Lentivirus generation and transduction

Lentiviral particles were generated according to standard procedures. In brief, HEK293T cells were co-transfected with the packaging vector psPAX2 (#12259, Addgene, Watertown, MA, USA), the envelope vector pMD2.G (#12260, Addgene) and the respective construct plasmid using TurboFect (Thermo Fisher Scientific, Waltham, MA, USA). After 48 hours, virus particle containing supernatant was collected and enriched using LentiConcentrator (OriGene). Cells were transduced with the respective concentrated virus particles using 10 µg/mL polybrene (Sigma-Aldrich) according to the manufacturer’s instructions. After 24 hours, medium was changed and cells cultivated for one week before starting with the respective experiments. Expression of the respective construct was confirmed via Western blot analysis.

### Generation and cultivation of murine keratinocytes

Murine keratinocytes were isolated from the epidermis of DSG2-W2A mut/mut and wt/wt mice and immortalized as described.^59^ In Brief, the epidermis of neonatal mice was separated from the dermis via incubation in dispase II solution (>2.4 U/ml dispase II, D4693, Sigma-Aldrich, in PBS) with 2x gentamicin/amphotericin B (CELLnTEC, Bern, Switzerland) over night at 4 °C. Keratinocytes were isolated by accutase (A6964, Sigma-Aldrich) for 20 minutes at room temperature and subsequent agitation. Released cells were cultured on collagen I coated plates (50201, IBIDI, Gräfelfing, Germany) in 0.06 mmol/l calcium murine keratinocyte medium (DMEM: Ham’s F12 3.5:1.1 mixture, SO-41660, PAN-Biotech, Aidenbach, Germany) supplemented with 10 % calcium-free foetal bovine serum (S0615, Merck), 2 mmol/l stable glutamine (BioConcept, Allschwil, Switzerland), 50μg/ml penicillin/streptomycin (Applichem), 10 ng/ml murine epidermal GF (Invitrogen, Carlsbad, CA), 1 mmol/l sodium pyruvate, 0.18 mmol/l adenine, 120 pmol/l cholera toxin, 5 µg/ml insulin, and 500 ng/ml hydrocortisone (all Sigma-Aldrich). Cells were kept in an incubator at 35 °C with 5 % CO_2_ and full humidity, the medium was changed every third day. When reaching confluency, cells were transferred into a new coated culture dish. After around six passages, cells were immortalized and could be expanded and seeded for experiments. 48 hours before experiments were conducted, 1.8 mmol/l calcium was added to the medium to induce cell differentiation. Cells were quarterly checked for mycoplasma infections using PCR and were proven negative.

### Cardiac slice culture

Mice were sacrificed by i.p. injection of pentobarbital. The heart was dissected by lateral thoracotomy and placed in ice-cold oxygenated slicing buffer (136 mmol/l NaCl, 5.4 mmol/l KCl, 1 mmol/l MgHPO_4_, 0.9 mmol/l CaCl_2_, 30 mmol/l 2,3-Butanedione monoxime, 5 mmol/l HEPES, 10 mmol/l glucose). After removal of both atria, the heart was embedded in 37 °C low melt agarose (Carl Roth, Karlsruhe, Germany) dissolved in slicing buffer without glucose. Using a LeicaVT1200 vibratome (Leica Biosystems, Nussloch, Germany), 300 μm thick tissue sections were cut with 1 mm amplitude and 0.07 mm/s speed. Freshly cut sections were transferred into ice-cold slicing buffer. For incubations, sections were transferred to 0.4 μm polycarbonate membranes cell culture inserts (VWR, Radnor, PA, USA) and incubated in Claycomb medium supplemented with 10 % foetal calf serum, 2 mmol/l L-glutamine, 10U/l : 10μg/ml penicillin and streptomycin (all from Sigma-Aldrich) at 37 °C, 5 % CO2 and treated with either rabbit anti-ITGαV/β6 1:15 (10D5, MAB2077Z, Sigma-Aldrich) with same amount of normal rabbit IgG (2729, Cell Signaling Technology, Danvers, MA, USA) as control IgG, or the selective TGFβ type I receptor inhibitor GW788388, 10 µmol/l (SML0116, Sigma-Aldrich, solved in DMSO) with DMSO as vehicle control. After treatment for 24 hours, cardiac slices were washed in HBSS on inserts and processed further for immunostaining, RNA isolation and Western blot analysis as described.

### Dissociation assay

Cells were treated as indicated and grown to confluency in 24-well plates. Cell monolayers were washed with HBSS and incubated with dissociation buffer (dispase II 2.5 U/mL, Sigma-Aldrich, D4693 in HBSS) at 37 °C till detachment of the cell monolayer from well bottom. After detachment, monolayers were mechanically stressed by defined pipetting using an electrical pipette (Eppendorf, Hamburg, Germany). The total number of resulting fragments per well was determined using a binocular stereo microscope (SZX2, Olympus). Fragments were counted if they were clearly visible at 1.25-fold magnification. The number of fragments is an indirect measure for intercellular cohesion. Images were acquired with a SLR camera (EOS 800D, Canon).

### Western blot analysis

Western blot analysis was performed using standard procedures. Tissue samples were homogenized in SDS-lysis buffer (12.5 mmol/l HEPES, 1 mmol/l EDTA, 12.5 mmol/l sodium fluoride, 0.5 % sodium dodecyl sulfate, pH 7.6) supplemented with protease inhibitor cocktail (cOmplete) and phosphatase inhibitor cocktail (PhosSTOP, both Roche, Basel, Switzerland) using FastPrep-24 5G bead beating grinder (MP Biomedicals, Santa Ana, CA, USA) and subsequently cleared by centrifugation. Confluent cell monolayers were washed with PBS and scraped in supplemented SDS lysis buffer. Lysates were sonicated and the total protein amount was determined with a BCA protein assay kit (Thermo Fisher Scientific) according to the manufacturer’s instructions. After lysates were denaturized for 5 minutes at 95 °C in Lämmli buffer, gel electrophoresis and wet blotting on nitrocellulose membranes (Novex, Thermo Fisher Scientific) were performed according to standard procedures. After a drying step, membranes were blocked in Intercept blocking buffer (Li-Cor, Lincoln, NE, USA) diluted 1:1 in TBS for 1 hour at room temperature. The following primary antibodies were incubated in antibody buffer (Intercept blocking buffer diluted 1:1 in TBS containing 0.2 % tween 20) at 4 °C overnight: Mouse anti-DSG1/2 (61002, Progen, Heidelberg, Germany), mouse anti-DSP (61003, Progen), mouse anti-PG (61005, Progen), mouse anti-N-Cadherin (NCAD, 610921, BD Bioscience, Franklin Lakes, NJ, USA), mouse anti-PKP2 (651101, Progen), rabbit anti-ITGB6 (ab187155, Abcam, Cambridge, UK), mouse anti-β-catenin (BCAT, 610154, BD Bioscience), rabbit anti-GAPDH (10494-1-AP, Proteintech, Rosemont, IL, USA), mouse anti-α-tubulin (ab7291, Abcam).

The secondary antibodies goat anti-mouse 800CW (925-32210) and goat anti-rabbit 680RD (925-68071, both Li-Cor) were incubated in TBS containing 0.1 % tween20 for 1 hour. Odyssey FC imaging system was used for fluorescence based band detection. Median band density was quantified applying ImageStudio (both Li-Cor) according to manufacturer’s instructions and normalized to the respective loading control.

### Immunostaining

For cryosections, tissue was embedded in 12 % mowiol 4-88, 5 % sorbitol, 0.5 % bovine serum albumin, 0.025 % sodium azide and frozen at −50 °C. 10 µm thick section were cut by a Cryo Star NX70 cryostat (Thermo Fisher Scientific), transferred to SuperFrost plus glass slides (Thermo Fisher Scientific), and air-dried. For immunostaining, sections were dried at 37 °C for 30 min, fixed in 2 % paraformaldehyde in PBS for 10 min, permeabilized with 0.2 % triton X-100 in PBS for 1 hour, and blocked with 3 % bovine serum albumin/0.12 % normal goat serum in PBS for 1 hour.

Fixed cardiac tissue embedded in paraffin was cut into 5 µm thick sections by an automated microtome (HM355S, Thermo Fisher Scientific). After deparaffinization, temperature-mediated antigen retrieval was performed in Tris/EGTA buffer (10 mmol/l Tris, 1 mmol/l EGTA, 0.05 % tween 20, pH 9) for 20 min at 95 °C. Tissue was permeabilized in 0.1 % triton X-100 in PBS for 5 min and blocked with 3 % bovine serum albumin/0.12 % normal goat serum in PBS for 1 hour.

The following primary antibodies were incubated in PBS at 4 °C overnight: Mouse anti-DSG1/2 (61002, Progen), mouse anti-DSP (61003, Progen), mouse anti-PG (61005, Progen), mouse anti-DSC2/3 (326200, Thermo Fisher Scientific), mouse anti-N-Cadherin (NCAD, 610921, BD Bioscience), rabbit anti-ITGB6 (ab187155, Abcam), rabbit anti-ITGB6/AV (BS-5791R, Bioss, Woburn, MA, USA), rabbit anti-ITGB1 (GTX112971, GeneTex, Irvine, CA, USA), rabbit anti-pSMAD2(S465/S467)/pSMAD3(S423/S425) (AP0548, Abclonal, Wuhan, China). Respective secondary goat anti-rabbit or goat anti-mouse antibodies coupled to Alexa Fluor 488, Alexa Fluor 568 (both Thermo Fisher Scientific), or cy5 (Dianova, Hamburg, Germany) were incubated for 1 hour at room temperature and DAPI (Sigma-Aldrich) was added for 10 minutes to counterstain nuclei. To visualize F-actin, phalloidin coupled to Dylight 488 (#21833, Thermo Fisher Scientific) or CruzFluor 647 (sc-363797, Santa Cruz Biotechnology, Dallas, TX, USA) was used. Tissue samples were mounted with Fluoromount Aqueous Mounting Medium (Sigma-Aldrich). For wide field image acquisition a 40x objective mounted on a Nanozoomer S60 slide scanner (Hamamatsu Photonics K.K., Hamamatsu, Japan) and for confocal image acquisition a 63x PL APO NA = 1.4 objective mounted on a LSM710 confocal microscope (Carl Zeiss, Jena, Germany) or a 63x HCX Plan-Apo NA = 1.4 objective mounted on a Leica SP5 confocal microscope (Leica Biosystems) was used. Fluorescence image analysis was performed by Fiji/ImageJ (NIH) or QuPath (QuPath developers, The University of Edinburgh, UK, version 0.2.3). For analysis of staining intensity in cardiomyocytes, a mask was generated using the f-acting signal and applied to the respective channel of interest to measure mean staining intensity in selected areas. For analysis of signal intensity in the nucleus, nuclei were detected via DAPI staining applying the QuPath cell detection tool. Respective masks were applied to the channel of interest to measure mean nuclear intensity in selected areas. For analysis of signal intensity at the ICDs, area masks of the ICDs were created and applied to the corresponding channels to measure mean signal intensity in these areas. The corresponding mean signal background was subtracted from all values. For lateral membranes, analysis mask was generated with a constant brush tool in QuPath and mean intensity measured.

### Histological staining

Tissue was embedded and cut as described in the section *Immunostaining*. Haematoxylin/ Eosin (HE) staining was performed according to standard procedures. In brief, sections were stained with Mayer’s haemalaun solution (Sigma-Aldrich) for 5 min, washed, dehydrated in an increasing ethanol series and stained with 0.5 % (w/v) eosin solution for 5 min. After washing steps in ethanol and methyl salicylate, sections were mounted with DPX Mountant (Sigma-Aldrich).

For Masson’s trichrome fibrosis, staining sections were fixed in 2 % paraformaldehyde in PBS for 10 min. After three washings in PBS and one rinse in distilled water, sections were re-fixed in Bouin’s solution (71 % picric acid, 24 % formaldehyde 37-40 %, 5 % glacial acetic acid) for 1 hour at 56 °C. Sections were then stained in Weigert’s iron haematoxylin solution (1:1 mixture of stock solution A [1 % haematoxylin in ethanol 95 %] and stock solution B [1.16 % ferric chloride, 1 % concentrated hydrochloric acid]) for 10 min and washed with water. A Biebrich scarlet-acid fuchsin solution (0.9 % Biebrich scarlet, 0.1 % acid fuchsin, 1 % glacial acetic acid) was applied as second staining for 13 min. Differentiation was performed in 5 % phosphotungstic acid for 20 min. Sections were transferred to aniline blue solution (2.5 % aniline blue, 2 % glacial acetic acid) and stained for 13 min. After a brief rinse in distilled water and 2 min differentiation in 1 % acetic acid, samples were dehydrated through 95 % ethanol and 100 % ethanol for 20 s each. Finally, sections were cleared in methyl salicylate and mounted in DPX Mountant (Sigma-Aldrich).

Images of histological sections were acquired with a 40x objective mounted on a Nanozoomer S60 slide scanner (Hamamatsu Photonics K.K.) and visualized with the NDP.view software (version 2.7.43, Hamamatsu Photonics K.K.). Area of fibrosis was analysed with the HALO Image Analysis Platform (v2.1.7, IndicaLabs, Albuquerque, NM, USA). The areas of interest (i.e. right and left ventricles) were defined using the layers. Cardiomyocytes and fibrotic tissue were then categorized with the Tissue Classifier Add-on based on their staining color. The Area Quantification FL analysis module was applied for quantifying cardiomyocytes and fibrotic areas. For analysis of hypertrophy, the area of 20 cross-sectioned cardiomyocytes with a central nucleus was measured for every heart using QuPath software (QuPath developers).

### Protein purification

For protein expression, the Fc control construct or the extracellular domain of murine Dsg2-WT or Dsg2-W2A were cloned into the Fc-His-pEGFP-N3 vector as described above. Wild type Chinese ovarian hamster cells (CHO) were transfected with the respective plasmids using TurboFect (Thermo Fisher Scientific) according to manufacturer’s instructions and selected with geneticin (VWR) for two weeks. After stable cells were grown to confluency, the cell culture supernatant was collected, the proteinase inhibitors leupeptin, aprotinin, pepstatin and phenylmethylsulfonylfluoride (all VWR) added and remaining cells removed by centrifugation and filtration. The cleared supernatant was added to a column containing HisLink Protein Purification Resin (V8823, Promega) and his-tag fusion proteins were purified according to manufacturer’s instructions. After imidazole-mediated elution, proteins were concentrated using Amicon Ultra-4 30 kDa centrifugal filter tubes (#UFC803024, Merck) and resuspended in sterile HBSS containing 1.2 mmol/l Ca^2+^. Protein concentration was determined by BCA protein assay kit (Thermo Fisher Scientific) and purity confirmed by Western blot analysis as described above.

### Single molecule force spectroscopy

For force spectroscopy experiments, a Nanowizard IV atomic force microscope (AFM, JPK Instruments, Berlin, Germany) mounted on an inverted fluorescence microscope (IX83, Olympus) was used. Recombinant proteins were generated as described in the section *Protein purification*. Flexible Si3N4 AFM probes (MLCT cantilever, Bruker, Billerica, MA, USA) and mica surfaces (Grade V-4, 01874-CA, Structure Probe, Inc., West Chester, PA, USA) were coated with aldehyde-PEG_20_-NHS ester spacer (BP-24296, Broadpharm, San Diego, CA, USA) to link recombinant molecules at a concentration of 0.15 mg/mL as described in ^60^. Force spectroscopy measurements were performed with the pyramid-shaped D-tip (nominal spring constant: 0.03 N/m) on functionalized mica sheets in HBSS containing 1.2 mmol/l Ca^2+^ and 0.1 % BSA at 37 °C. Spring constant was calibrated for each cantilever at 37 °C applying the thermal noise method.^61^ Force spectroscopy experiments were performed in force mapping mode using following settings: relative setpoint 0.4 nN, z-length 0.3 – 0.5 µm, extend delay 0.1 s, pulling speed as indicated ranging from 0.5 µm/s to 15 µm/s, scanning area: 10 µm x 10 µm, 25 px x 25 px and recorded with the SPM Control v.4 software (JPK Instruments). Force distance blots were analysed using JPKSPM Data Processing software (version 6, JPK Instruments). For calculation of force histogram, extreme peak curve fit and application of Bell’s equation, Origin software (Originlab, Northampton, MA, USA) was used.

### Echocardiography with Electrocardiogram (ECG)

Transthoracic echocardiography was performed using the Vevo 2100 ultrasound system (VisualSonics, Toronto, ON Canada) equipped with a MS-550 linear-array probe working at a central frequency of 40 MHz. After the animals were anesthetized with 3.0 % (v/v) isoflurane carried by pure oxygen, they were placed at supine position on a pre-warmed imaging platform. Anesthesia was maintained by 1.5 % (v/v) isoflurane through a nose cone and the body temperature was controlled at around 37 °C by a rectal thermocouple probe. Eye gel (Lacrinorm) was applied to prevent ocular dehydration. Needle probes attached to ECG leads embedded in the imaging platform were inserted subcutaneously to each limb for ECG recording. Hairs on the chest were removed by applying commercially available hair removal cream (Nair). Cardiac geometry and function were evaluated using 2-D guided M-mode at the mid-papillary muscle level from parasternal short-axis view before and 2, 4, 10, 11, 12 and 13 min after injection of isoprenaline hydrochloride (I5627, Sigma-Aldrich) at 2 mg/kg body weight i.p.. ECG was monitored full time during the whole procedure and arrhythmia was recorded and counted. Data was transferred to an offline computer and analyzed with Vevo 2100 software (version 1.6.0, VisualSonics) by an investigator blinded to the study groups. Left ventricular anterior (LVAW) and posterior (LVPW) wall thickness and internal dimensions (LVID) were measured from the M-mode during systole (s) and diastole (d). Values were averages of three cardiac cycles. Left ventricular ejection fraction (EF) was calculated from derived volumes (Vol), which are computed based on the Teichholz formula (LV Vol;d = (7.0 / (2.4 + LVID;d)) × LVID;d^3^, LV Vol;s = (7.0 / (2.4 + LVID;s)) × LVID;s^3^, EF % = 100 × ((LV Vol;d – LV Vol;s) / LV Vol;d)). Left ventricular mass (LV Mass) was calculated based on a corrected cube model (LV Mass = 1.053 × ((LVAW;d + LVID;d + LVPW;d)^3^ – LVID;d^3^) × 0.8). After echocardiography, mice were euthanized via cervical dislocation and hearts were dissected. Heart and body weight was determined and organs were embedded and stained as described in the section *Histological staining* for further analysis to determine the amount of fibrosis.

### Transmission electron microscopy (TEM)

Ventricular cardiac tissue was dissected, cut and fixed in 2 % paraformaldehyde and 2.5 % glutaraldehyde (Electron Microscopy Sciences, Hatfield, PA, USA) in 0.1 mol/l Pipes, 2 mmol/l CaCl_2_, pH 7.3. After 15 min incubation, fixative was renewed and incubated at 4 °C for 16 hours. Samples were washed three times with cold 0.1 mol/l Pipes, 2 mmol/l CaCl_2_, pH 7.3 and rinsed with 0.1 mol/l cacodylate buffer, pH 7.3. Post-fixation was performed for 1 hour at 4 °C using 1 % osmium tetroxide and 0.8 % potassium ferracyanide (Electron Microscopy Sciences) in 0.1 mol/l cacodylate buffer, pH 7.3. After washing steps with cacodylate buffer, pH 7.3 and ultrapure distilled water, tissue samples were stained with 1 % aqueous uranyl acetate (Electron Microscopy Sciences) for 1 hour at 4 °C. Dehydration was performed by an ascending ethanol series at 4 °C. After three washes with 100 % ethanol, samples were rinsed in acetone and first embedded in a mixture of resin/acetone followed by pure Epon 812 resin (Electron Microscopy Sciences) over night. Samples were mounted on BEEM capsules (Electron Microscopy Sciences) filled with EPON. After polymerization at 60 °C for 48 hours, samples were removed from the EPON block with the nitrogen hot water method. 70 nm thin serial sections, cut with a diamond knife, were mounted on formvar-carbon coated copper slot grids, stained with uranyl acetate and Reynolds’s lead citrate. Samples were examined in a FEI Tecnai T12 spirit transmission electron microscope (Thermo Fisher Scientific) operating at 80 kV equipped with a CCD Veleta digital camera.

### RNA isolation

Cardiac tissue was washed in ice-cold HBSS and lysed in TRI reagent (Molecular Research Center, Inc., Cincinnati, OH, USA). Tissue homogenization was conducted via the FastPrep-24 5G bead beating grinder (MP Biomedicals, Santa Ana, CA, USA) using 2.8 mm stainless steel beads (Sigma-Aldrich) according to manufacturer’s protocol with subsequent centrifugation to clear the lysate. RNA was isolated via the Direct-zol RNA MiniPrep kit including Zymo-Spin II and DNAse restriction step (R2050, Zymo research, Irvine, CA, USA).

### Quantitative Real time PCR (q-RT-PCR)

RNA was isolated as described above. Quantity and quality of RNA was determined by Nanodrop 1000 Spectrophotometer (Thermo Fisher Scientific). Up to 1 µg of isolated RNA was used for reverse transcription with SuperScript III (Thermo Fisher Scientific). Quantitative real time PCR was performed with StepOne Real time PCR Systems (Applied Biosystems) using Power SYBR Green PCR Master Mix (Thermo Fisher Scientific). Primers are listed in Supplementary Table 1. As reference, the mean Ct value of *Gapdh* and *Tubg2* of the respective sample was used.

### Mouse RNA sequencing (RNA-Seq)

For transcriptomic analysis before and after onset of fibrosis, hearts of 5-days- and 9-weeks-old mice were dissected. Wt/wt and mut/mut mice were matched for age and sex. For 5-days-old mice, atria were removed and both ventricles lysed. For 9-weeks-old animals, similar sized tissue samples were taken from the right and left ventricle via a 3 mm diameter biopsy punch (Viollier, Allschwil, Switzerland). RNA was isolated as described in the section *RNA isolation*. RNA were quality-checked on the TapeStation instrument (Agilent Technologies, Santa Clara, CA, USA) using the RNA ScreenTape (Agilent, #5067-5576). RNA were quantified by Fluorometry using the QuantiFluor RNA System (#E3310, Promega). Library preparation was performed, starting from 200ng total RNA, using the TruSeq Stranded mRNA Library Kit (#20020595, Illumina, San Diego, CA, USA) and the TruSeq RNA UD Indexes (#20022371, Illumina). 15 cycles of PCR were performed. Libraries were quality-checked on the Fragment Analyzer (Advanced Analytical, Ames, IA, USA) using the Standard Sensitivity NGS Fragment Analysis Kit (#DNF-473, Advanced Analytical) revealing excellent quality of libraries (average concentration was 179±9 nmol/l and average library size was 329±3 base pairs). For Sequencing, samples were pooled to equal molarity. The pool was quantified by Fluorometry using the QuantiFluor ONE dsDNA System (#E4871, Promega). Libraries were sequenced Paired-End 38 bases (in addition: 8 bases for index 1 and 8 bases for index 2) using the NextSeq 500 High Output Kit 75-cycles (#FC-404-1005, Illumina) loaded at 1.8pM and including 1 % PhiX. Primary data analysis was performed with the Illumina RTA version 2.11.3. On average per sample: 38.7±4.6 million pass-filter reads were collected on that NextSeq 500 Flow-Cell.

### Mouse RNA-Seq data analysis

Reads were aligned to the mouse mm10 genome using the aligner STAR (version 2.7.3a) ^62^ with extra options “--outFilterMultimapNmax 10 --outSAMmultNmax 1” for handling multimapping reads. Aligned reads were assigned to ensembl genes (version 101) using the tool featureCounts from the subread package (version 2.0.1) ^63^ with extra options “-O -M --read2pos5 –primary-s 2”. All further analysis steps were performed with R/Bioconductor (R version 4.0.3, Bioconductor version 2.50.0). Gene counts were loaded into R and differential gene expression analysis followed the edgeR workflow ^64^. Specifically, genes were filtered for expression using the function filterByExpr which retained 19504 genes. Samples were classified into 4 groups according to genotype (wt/wt and mut/mut) and time point (5 days and 9 weeks) and differential gene expression was performed between all 4 groups using the functions glmQLFit and glmQLFTest. Gene set enrichment analysis was performed using the collections of gene sets form MSigDb (https://www.gseamsigdb.org/gsea/msigdb/collections.jsp) or selected gene sets as indicated and relied on the edgeR function camera. External gene lists were tested on different contrasts using the cameraPR function of edgeR.

### Re-analysis of available ACM data sets and comparison with murine data sets

Raw data of GEO data sets GSE107157 and GSE107480 were downloaded from the European Nucleotide Archive and mapped to the human hg38AnalysisSet genome using STAR. Gene expression was quantified with featureCounts and relied on the human ensembl gene annotation (version 96). Gene counts of both data sets were imported into R and analysed using the edgeR workflow. Specifically, genes were filtered by expression which retained 25293 genes, followed by a differential expression analysis contrasting the 4 sample groups which arise from tissue source (left and right ventricle) and disease (ARVC and healthy). In order to compare mouse specific gene lists to the human data, mouse genes were mapped to human orthologs using BioMart. The gene set of human orthologs was tested for differential enrichment in the mouse gene expression contrasts defined above using the function cameraPR.

### Immunoprecipitation (IP)

Tissue samples were lysed in modified RIPA buffer (10 mmol/l Na_2_HPO_4_, 150 mmol/l NaCl, 1 % triton X-100, 0.25 % SDS, 1 % sodium deoxycholate, pH 7.3) complemented with aprotinin, leupeptin, pepstatin, phenylmethylsulfonyl fluoride and PhosSTOP by homogenization via the FastPrep-24 5G bead beating grinder (MP Biomedicals) using 2.8 mm stainless steel beads (Sigma-Aldrich). Samples were centrifuged at 12000 G, 5 min at 4 °C and cleared supernatant used for further analysis. Protein concentration was determined using the BCA method (Thermo Fisher Scientific). Equal amounts of protein were incubated with rabbit anti-ITGB6 (ab187155, Abcam), or normal rabbit IgG (#sc-2027, Santa Cruz) overnight at 4 °C. Lysates were incubated with pre-washed, magnetic Dynabeads Protein G (Thermo Fisher Scientific) for 1 hour at 4 °C on a rotor. Beads were washed 10 times and denaturized at 95 °C in Lämmli buffer for 10 minutes. The samples were loaded on SDS polyacrylamide gels and Western blot analysis was performed as described above.

### Structured illumination microscopy (SIM)

For structured illumination microscopy (SIM) of cryopreserved mouse heart tissue, sectioning and immunostaining were performed as described. Image stacks spanning full ICDs were acquired using a DeltaVision OMX-Blaze (Version 4; Applied Precision) equipped with a 60x PL APO NA = 1.42 objective (Olympus) yielding a voxel size of 0.04 x 0.04 x 0.125 µm.

For analysis of DSG2 and ITGB6 particle size and frequency within ICDs, the image stacks (typically containing one clearly defined ICD) were loaded into Fiji. The ICD was outlined as ROI in a maximum intensity projection, which was also used to determine the ICD volume in the stack. DSG2 and ITGB6 signals within the ICD were thresholded and annotated as objects using the 3D Object Counter tool.^65^ Annotated objects were further analyzed with regard to volume and number using the 3D Manager from the 3D Suite plugin.^66^ The Imaris software package (v. 9.6, Oxford Instruments, Zurich, Switzerland) was used for 3D image reconstruction and for analysis of ITGB6 distribution and spatial proximity between DSG2 and ITGB6. ICDs were first isolated by manually creating a surface containing the entire ICD. ITGB6 and DSG2 signals were then automatically detected and annotated as individual objects using similar thresholding values. The Statistics module was applied to calculate spatial relationships (average distance to nearest neighbor objects of the same channel, distance to the nearest neighbor of the respective other channel).

### Statistics and data compilation

Figures were compiled with Adobe Photoshop CC 2017 and Adobe Illustrator CC 2017 (both Adobe, San José, CA). Statistical computations were performed with Prism 8 (GraphPad Software, La Jolla, CA). The statistical tests used to compare the respective data sets are mentioned in the corresponding figure legend. Statistical significance was assumed at *P* < 0.05. Unless otherwise stated, data are presented as dot blot, with each dot representing the mean of the respective technical replicates of one biological replicate. Each animal or independent seeding of cells were taken as biological replicate. The mean value of these dots is shown as bar diagram ± standard deviation.

## Acknowledgement

We are grateful to Alain Brühlhart and the animal care team from the Animal Facility (University of Basel, Switzerland), Dr. Cinzia Tiberi from the Center for Cellular Imaging and NanoAnalytics (C-CINA, University of Basel), Dr. Alexia Loynton-Ferrand from the Imaging Core Facility (IMCF, Biocenter, University of Basel), Dr. Diego Calabrese from the Histology Core Facility, Dr. Mike Abanto, Dr. Beat Erne, and Ewelina Bartoszek from the Microscopy Core Facility (all DBM, University of Basel), Dr. Christian Beisel and Philippe Demougin from the Genomics Facility Basel (D-BSSE, ETH Zürich, and University of Basel, Switzerland) for excellent support. We thank Nicolas Schlegel (Department of General, Visceral, Vascular and Pediatric Surgery, University Hospital Würzburg, Würzburg, Germany) for providing the CaCo2(DSG2 KO) cell line, and Nikola Golenhofen (Institute of Anatomy and Cell Biology, University of Ulm, Ulm, Germany) for providing the Fc-His-pEGFP-N3 plasmid. We thank Anja Fuchs for excellent technical assistance. Calculations were performed at sciCORE (http://scicore.unibas.ch/) scientific computing center at University of Basel.

## Data availability

RNA-Seq data have been deposited in the public database GEO accession number GSE181868. Additional data pertaining to the present article are available from the corresponding author upon request.

## References

1. Corrado D, Link MS and Calkins H. Arrhythmogenic Right Ventricular Cardiomyopathy. N Engl J Med. 2017;376:1489–90.

2. Austin KM, Trembley MA, Chandler SF, Sanders SP, Saffitz JE, Abrams DJ and Pu WT. Molecular mechanisms of arrhythmogenic cardiomyopathy. Nat Rev Cardiol. 2019;16:519–537.

3. Thiene G, Nava A, Corrado D, Rossi L and Pennelli N. Right ventricular cardiomyopathy and sudden death in young people. The New England journal of medicine. 1988;318:129–33.

4. Towbin JA, McKenna WJ, Abrams DJ, Ackerman MJ, Calkins H, Darrieux FCC, Daubert JP, de Chillou C, DePasquale EC, Desai MY, Estes NAM, 3rd, Hua W, Indik JH, Ingles J, James CA, John RM, Judge DP, Keegan R, Krahn AD, Link MS, Marcus FI, McLeod CJ, Mestroni L, Priori SG, Saffitz JE, Sanatani S, Shimizu W, van Tintelen JP, Wilde AAM and Zareba W. 2019 HRS expert consensus statement on evaluation, risk stratification, and management of arrhythmogenic cardiomyopathy. Heart Rhythm. 2019;16:e301–e372.

5. Delva E, Tucker DK and Kowalczyk AP. The desmosome. Cold Spring Harb Perspect Biol. 2009;1:a002543.

6. Delmar M. The intercalated disk as a single functional unit. Heart rhythm. 2004;1:12–3.

7. Delmar M and McKenna WJ. The cardiac desmosome and arrhythmogenic cardiomyopathies: from gene to disease. Circ Res. 2010;107:700–14.

8. Franke WW, Borrmann CM, Grund C and Pieperhoff S. The area composita of adhering junctions connecting heart muscle cells of vertebrates. I. Molecular definition in intercalated disks of cardiomyocytes by immunoelectron microscopy of desmosomal proteins. European journal of cell biology. 2006;85:69–82.

9. Leo-Macias A, Agullo-Pascual E and Delmar M. The cardiac connexome: Non-canonical functions of connexin43 and their role in cardiac arrhythmias. Seminars in cell & developmental biology. 2016;50:13–21.

10. Basso C, Czarnowska E, Della Barbera M, Bauce B, Beffagna G, Wlodarska EK, Pilichou K, Ramondo A, Lorenzon A, Wozniek O, Corrado D, Daliento L, Danieli GA, Valente M, Nava A, Thiene G and Rampazzo A. Ultrastructural evidence of intercalated disc remodelling in arrhythmogenic right ventricular cardiomyopathy: an electron microscopy investigation on endomyocardial biopsies. Eur Heart J. 2006;27:1847–54.

11. Zorzi A, Cipriani A, Mattesi G, Vio R, Bettella N and Corrado D. Arrhythmogenic Cardiomyopathy and Sports Activity. J Cardiovasc Transl Res. 2020;13:274–283.

12. Kirchhof P, Fabritz L, Zwiener M, Witt H, Schafers M, Zellerhoff S, Paul M, Athai T, Hiller KH, Baba HA, Breithardt G, Ruiz P, Wichter T and Levkau B. Age- and training-dependent development of arrhythmogenic right ventricular cardiomyopathy in heterozygous plakoglobin-deficient mice. Circulation. 2006;114:1799–806.

13. Hariharan V, Asimaki A, Michaelson JE, Plovie E, MacRae CA, Saffitz JE and Huang H. Arrhythmogenic right ventricular cardiomyopathy mutations alter shear response without changes in cell-cell adhesion. Cardiovascular research. 2014;104:280–9.

14. Huang H, Asimaki A, Lo D, McKenna W and Saffitz J. Disparate effects of different mutations in plakoglobin on cell mechanical behavior. Cell Motil Cytoskeleton. 2008;65:964–78.

15. Schlipp A, Schinner C, Spindler V, Vielmuth F, Gehmlich K, Syrris P, McKenna WJ, Dendorfer A, Hartlieb E and Waschke J. Desmoglein-2 interaction is crucial for cardiomyocyte cohesion and function. Cardiovasc Res. 2014;104:245–57.

16. Sato PY, Coombs W, Lin X, Nekrasova O, Green KJ, Isom LL, Taffet SM and Delmar M. Interactions between ankyrin-G, Plakophilin-2, and Connexin43 at the cardiac intercalated disc. Circulation research. 2011;109:193–201.

17. Corrado D, Basso C and Judge DP. Arrhythmogenic Cardiomyopathy. Circ Res. 2017;121:784–802.

18. Gerull B and Brodehl A. Genetic Animal Models for Arrhythmogenic Cardiomyopathy. Front Physiol. 2020;11:624.

19. Harrison OJ, Brasch J, Lasso G, Katsamba PS, Ahlsen G, Honig B and Shapiro L. Structural basis of adhesive binding by desmocollins and desmogleins. Proc Natl Acad Sci U S A. 2016;113:7160–5.

20. Al-Amoudi A and Frangakis AS. Structural studies on desmosomes. Biochem Soc Trans. 2008;36:181–7.

21. Bell GI. Models for the specific adhesion of cells to cells. Science. 1978;200:618–27.

22. Vielmuth F, Hartlieb E, Kugelmann D, Waschke J and Spindler V. Atomic force microscopy identifies regions of distinct desmoglein 3 adhesive properties on living keratinocytes. Nanomedicine. 2015;11:511–20.

23. Baumgartner W, Hinterdorfer P, Ness W, Raab A, Vestweber D, Schindler H and Drenckhahn D. Cadherin interaction probed by atomic force microscopy. Proc Natl Acad Sci U S A. 2000;97:4005–10.

24. Waschke J, Menendez-Castro C, Bruggeman P, Koob R, Amagai M, Gruber HJ, Drenckhahn D and Baumgartner W. Imaging and force spectroscopy on desmoglein 1 using atomic force microscopy reveal multivalent Ca(2+)- dependent, low-affinity trans-interaction. J Membr Biol. 2007;216:83–92.

25. Meir M, Burkard N, Ungewiss H, Diefenbacher M, Flemming S, Kannapin F, Germer CT, Schweinlin M, Metzger M, Waschke J and Schlegel N. Neurotrophic factor GDNF regulates intestinal barrier function in inflammatory bowel disease. J Clin Invest. 2019;129:2824–2840.

26. Ishii K, Harada R, Matsuo I, Shirakata Y, Hashimoto K and Amagai M. In vitro keratinocyte dissociation assay for evaluation of the pathogenicity of anti-desmoglein 3 IgG autoantibodies in pemphigus vulgaris. J Invest Dermatol. 2005;124:939–46.

27. Corrado D, Perazzolo Marra M, Zorzi A, Beffagna G, Cipriani A, Lazzari M, Migliore F, Pilichou K, Rampazzo A, Rigato I, Rizzo S, Thiene G, Anastasakis A, Asimaki A, Bucciarelli-Ducci C, Haugaa KH, Marchlinski FE, Mazzanti A, McKenna WJ, Pantazis A, Pelliccia A, Schmied C, Sharma S, Wichter T, Bauce B and Basso C. Diagnosis of arrhythmogenic cardiomyopathy: The Padua criteria. Int J Cardiol. 2020;319:106–114.

28. Gaertner A, Schwientek P, Ellinghaus P, Summer H, Golz S, Kassner A, Schulz U, Gummert J and Milting H. Myocardial transcriptome analysis of human arrhythmogenic right ventricular cardiomyopathy. Physiol Genomics. 2012;44:99–109.

29. Broussard JA, Getsios S and Green KJ. Desmosome regulation and signaling in disease. Cell Tissue Res. 2015;360:501–12.

30. Spindler V and Waschke J. Desmosomal cadherins and signaling: lessons from autoimmune disease. Cell Commun Adhes. 2014;21:77–84.

31. Margadant C and Sonnenberg A. Integrin-TGF-beta crosstalk in fibrosis, cancer and wound healing. EMBO Rep. 2010;11:97–105.

32. Munger JS and Sheppard D. Cross talk among TGF-beta signaling pathways, integrins, and the extracellular matrix. Cold Spring Harb Perspect Biol. 2011;3:a005017.

33. Koivisto L, Bi J, Hakkinen L and Larjava H. Integrin alphavbeta6: Structure, function and role in health and disease. Int J Biochem Cell Biol. 2018;99:186–196.

34. Han H, Cho JW, Lee S, Yun A, Kim H, Bae D, Yang S, Kim CY, Lee M, Kim E, Lee S, Kang B, Jeong D, Kim Y, Jeon HN, Jung H, Nam S, Chung M, Kim JH and Lee I. TRRUST v2: an expanded reference database of human and mouse transcriptional regulatory interactions. Nucleic Acids Res. 2018;46:D380–D386.

35. Bian Q, Ma L, Jain A, Crane JL, Kebaish K, Wan M, Zhang Z, Edward Guo X, Sponseller PD, Seguin CA, Riley LH, Wang Y and Cao X. Mechanosignaling activation of TGFbeta maintains intervertebral disc homeostasis. Bone Res. 2017;5:17008.

36. Ritsma L, Dey-Guha I, Talele N, Sole X, Salony, Chowdhury J, Ross KN and Ramaswamy S. Integrin beta1 activation induces an anti-melanoma host response. PLoS One. 2017;12:e0175300.

37. Gellibert F, de Gouville AC, Woolven J, Mathews N, Nguyen VL, Bertho-Ruault C, Patikis A, Grygielko ET, Laping NJ and Huet S. Discovery of 4-{4-[3-(pyridin-2-yl)-1H-pyrazol-4-yl]pyridin-2-yl}-N-(tetrahydro-2H-pyran-4-yl)benzamide (GW788388): a potent, selective, and orally active transforming growth factor-beta type I receptor inhibitor. J Med Chem. 2006;49:2210–21.

38. Medeiros-Domingo A, Saguner AM, Magyar I, Bahr A, Akdis D, Brunckhorst C, Duru F and Berger W. Arrhythmogenic right ventricular cardiomyopathy: implications of next-generation sequencing in appropriate diagnosis. Europace. 2017;19:1063–1069.

39. DeWitt ES, Chandler SF, Hylind RJ, Beausejour Ladouceur V, Blume ED, VanderPluym C, Powell AJ, Fynn-Thompson F, Roberts AE, Sanders SP, Bezzerides V, Lakdawala NK, MacRae CA and Abrams DJ. Phenotypic Manifestations of Arrhythmogenic Cardiomyopathy in Children and Adolescents. J Am Coll Cardiol. 2019;74:346–358.

40. Krusche CA, Holthofer B, Hofe V, van de Sandt AM, Eshkind L, Bockamp E, Merx MW, Kant S, Windoffer R and Leube RE. Desmoglein 2 mutant mice develop cardiac fibrosis and dilation. Basic Res Cardiol. 2011;106:617–33.

41. Wang Y, Li C, Shi L, Chen X, Cui C, Huang J, Chen B, Hall DD, Pan Z, Lu M, Hong J, Song LS and Zhao S. Integrin beta1D Deficiency-Mediated RyR2 Dysfunction Contributes to Catecholamine-Sensitive Ventricular Tachycardia in Arrhythmogenic Right Ventricular Cardiomyopathy. Circulation. 2020;141:1477–1493.

42. Puzzi L, Borin D, Gurha P, Lombardi R, Martinelli V, Weiss M, Andolfi L, Lazzarino M, Mestroni L, Marian AJ and Sbaizero O. Knock Down of Plakophillin 2 Dysregulates Adhesion Pathway through Upregulation of miR200b and Alters the Mechanical Properties in Cardiac Cells. Cells. 2019;8.

43. Meecham A and Marshall JF. The ITGB6 gene: its role in experimental and clinical biology. Gene X. 2020;5:100023.

44. Munger JS, Huang X, Kawakatsu H, Griffiths MJ, Dalton SL, Wu J, Pittet JF, Kaminski N, Garat C, Matthay MA, Rifkin DB and Sheppard D. The integrin alpha v beta 6 binds and activates latent TGF beta 1: a mechanism for regulating pulmonary inflammation and fibrosis. Cell. 1999;96:319–28.

45. Dutta A, Li J, Fedele C, Sayeed A, Singh A, Violette SM, Manes TD and Languino LR. alphavbeta6 integrin is required for TGFbeta1-mediated matrix metalloproteinase2 expression. Biochem J. 2015;466:525–36.

46. Hanna A, Humeres C and Frangogiannis NG. The role of Smad signaling cascades in cardiac fibrosis. Cell Signal. 2021;77:109826.

47. Frangogiannis NG. Cardiac fibrosis: Cell biological mechanisms, molecular pathways and therapeutic opportunities. Mol Aspects Med. 2019;65:70–99.

48. Li D, Liu Y, Maruyama M, Zhu W, Chen H, Zhang W, Reuter S, Lin SF, Haneline LS, Field LJ, Chen PS and Shou W. Restrictive loss of plakoglobin in cardiomyocytes leads to arrhythmogenic cardiomyopathy. Hum Mol Genet. 2011;20:4582–96.

49. Dubash AD, Kam CY, Aguado BA, Patel DM, Delmar M, Shea LD and Green KJ. Plakophilin-2 loss promotes TGF-beta1/p38 MAPK-dependent fibrotic gene expression in cardiomyocytes. J Cell Biol. 2016;212:425–38.

50. Chen C, Li R, Ross RS and Manso AM. Integrins and integrin-related proteins in cardiac fibrosis. J Mol Cell Cardiol. 2016;93:162–74.

51. Kulkarni AB, Huh CG, Becker D, Geiser A, Lyght M, Flanders KC, Roberts AB, Sporn MB, Ward JM and Karlsson S. Transforming growth factor beta 1 null mutation in mice causes excessive inflammatory response and early death. Proc Natl Acad Sci U S A. 1993;90:770–4.

52. Puthawala K, Hadjiangelis N, Jacoby SC, Bayongan E, Zhao Z, Yang Z, Devitt ML, Horan GS, Weinreb PH, Lukashev ME, Violette SM, Grant KS, Colarossi C, Formenti SC and Munger JS. Inhibition of integrin alpha(v)beta6, an activator of latent transforming growth factor-beta, prevents radiation-induced lung fibrosis. Am J Respir Crit Care Med. 2008;177:82–90.

53. John AE, Graves RH, Pun KT, Vitulli G, Forty EJ, Mercer PF, Morrell JL, Barrett JW, Rogers RF, Hafeji M, Bibby LI, Gower E, Morrison VS, Man Y, Roper JA, Luckett JC, Borthwick LA, Barksby BS, Burgoyne RA, Barnes R, Le J, Flint DJ, Pyne S, Habgood A, Organ LA, Joseph C, Edwards-Pritchard RC, Maher TM, Fisher AJ, Gudmann NS, Leeming DJ, Chambers RC, Lukey PT, Marshall RP, Macdonald SJF, Jenkins RG and Slack RJ. Translational pharmacology of an inhaled small molecule alphavbeta6 integrin inhibitor for idiopathic pulmonary fibrosis. Nat Commun. 2020;11:4659.

54. Patsenker E, Popov Y, Stickel F, Jonczyk A, Goodman SL and Schuppan D. Inhibition of integrin alphavbeta6 on cholangiocytes blocks transforming growth factor-beta activation and retards biliary fibrosis progression. Gastroenterology. 2008;135:660–70.

55. Popov Y, Patsenker E, Stickel F, Zaks J, Bhaskar KR, Niedobitek G, Kolb A, Friess H and Schuppan D. Integrin alphavbeta6 is a marker of the progression of biliary and portal liver fibrosis and a novel target for antifibrotic therapies. J Hepatol. 2008;48:453–64.

56. Concordet JP and Haeussler M. CRISPOR: intuitive guide selection for CRISPR/Cas9 genome editing experiments and screens. Nucleic Acids Res. 2018;46:W242–W245.

57. Chen S, Lee B, Lee AY, Modzelewski AJ and He L. Highly Efficient Mouse Genome Editing by CRISPR Ribonucleoprotein Electroporation of Zygotes. J Biol Chem. 2016;291:14457–67.

58. Haueter S, Kawasumi M, Asner I, Brykczynska U, Cinelli P, Moisyadi S, Burki K, Peters AH and Pelczar P. Genetic vasectomy-overexpression of Prm1-EGFP fusion protein in elongating spermatids causes dominant male sterility in mice. Genesis. 2010;48:151–60.

59. Hiermaier M, Kliewe F, Schinner C, Studle C, Maly IP, Wanuske MT, Rotzer V, Endlich N, Vielmuth F, Waschke J and Spindler V. The Actin-Binding Protein alpha-Adducin Modulates Desmosomal Turnover and Plasticity. J Invest Dermatol. 2021;141:1219–1229 e11.

60. Ebner A, Wildling L, Kamruzzahan AS, Rankl C, Wruss J, Hahn CD, Holzl M, Zhu R, Kienberger F, Blaas D, Hinterdorfer P and Gruber HJ. A new, simple method for linking of antibodies to atomic force microscopy tips. Bioconjug Chem. 2007;18:1176–84.

61. Hutter JL and Bechhoefer J. Calibration of atomic-force microscope tips. Review of Scientific Instruments. 1993;64:1868–1873.

62. Dobin A, Davis CA, Schlesinger F, Drenkow J, Zaleski C, Jha S, Batut P, Chaisson M and Gingeras TR. STAR: ultrafast universal RNA-seq aligner. Bioinformatics. 2013;29:15–21.

63. Liao Y, Smyth GK and Shi W. featureCounts: an efficient general purpose program for assigning sequence reads to genomic features. Bioinformatics. 2014;30:923–30.

64. Chopin M, Preston SP, Lun ATL, Tellier J, Smyth GK, Pellegrini M, Belz GT, Corcoran LM, Visvader JE, Wu L and Nutt SL. RUNX2 Mediates Plasmacytoid Dendritic Cell Egress from the Bone Marrow and Controls Viral Immunity. Cell Rep. 2016;15:866–878.

65. Bolte S and Cordelieres FP. A guided tour into subcellular colocalization analysis in light microscopy. J Microsc. 2006;224:213–32.

66. Ollion J, Cochennec J, Loll F, Escude C and Boudier T. TANGO: a generic tool for high-throughput 3D image analysis for studying nuclear organization. Bioinformatics. 2013;29:1840–1.

67. Liberzon A, Subramanian A, Pinchback R, Thorvaldsdottir H, Tamayo P and Mesirov JP. Molecular signatures database (MSigDB) 3.0. Bioinformatics. 2011;27:1739–40.

