## Supplemental Material for "Defective Desmosomal Adhesion Causes Arrhythmogenic Cardiomyopathy by involving an Integrin-αVβ6/TGF-β Signaling Cascade"

### Supplementary Figures and Figure Legends

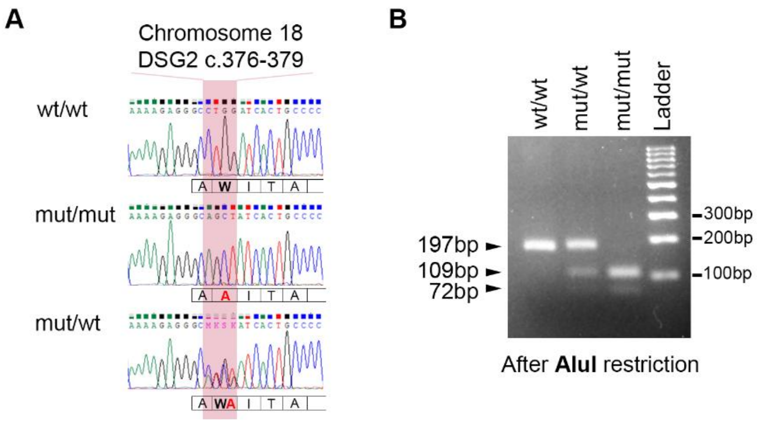


**Supplementary Figure 1. Sequencing and genotyping of the DSG2-W2A mouse model.** (**A**) Representative sequencing results of the DSG2-W2A mouse model for the indicated genotypes. The bars below show the related translation in amino acids. (**B**) Representative results of DSG2-W2A genotyping showing electrophoresis after restriction of the PCR product with AluI.

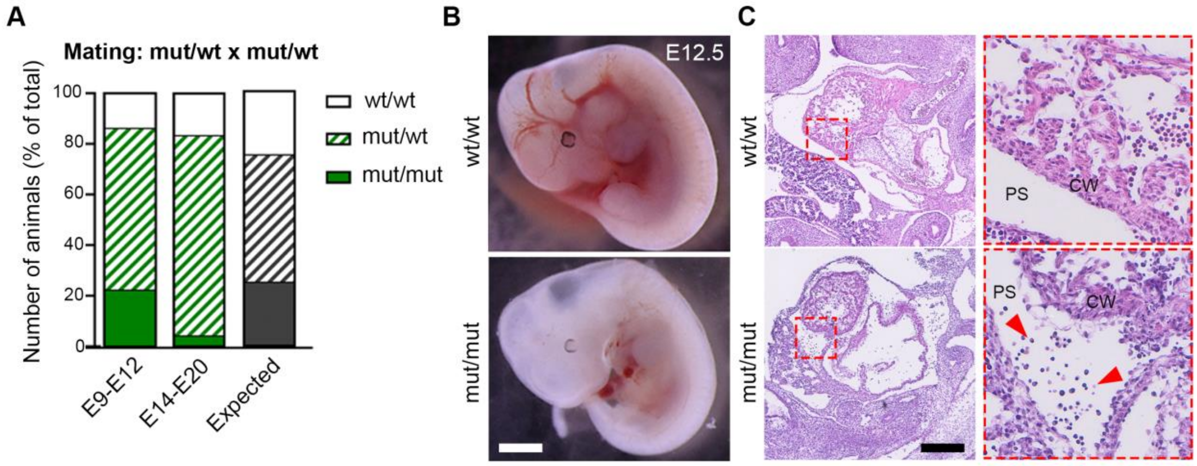


**Supplementary Figure 2. Loss of mut/mut animals during development due to pericardial bleeding.** (**A**) Genotype analysis of embryos derived from matings of mut/wt animals between developmental day E9 to E20 compared to the expected genotype distribution according to Mendel in gray. (**B**) Macroscopic appearance of viable embryos at day E12.5. Scale bar: 2 mm. (**C**) Haematoxylin/Eosin staining of sagittal sections of the cardiac area derived from embryos in **B**. Red rectangle depicts area of zoomed insert. Scale bar overview: 1 mm. Red arrowheads point to blood precursor cells in the pericardial space (PS) adjacent to the cardiac wall (CW). Images are representative for 4 embryos per genotype from 4 litters.

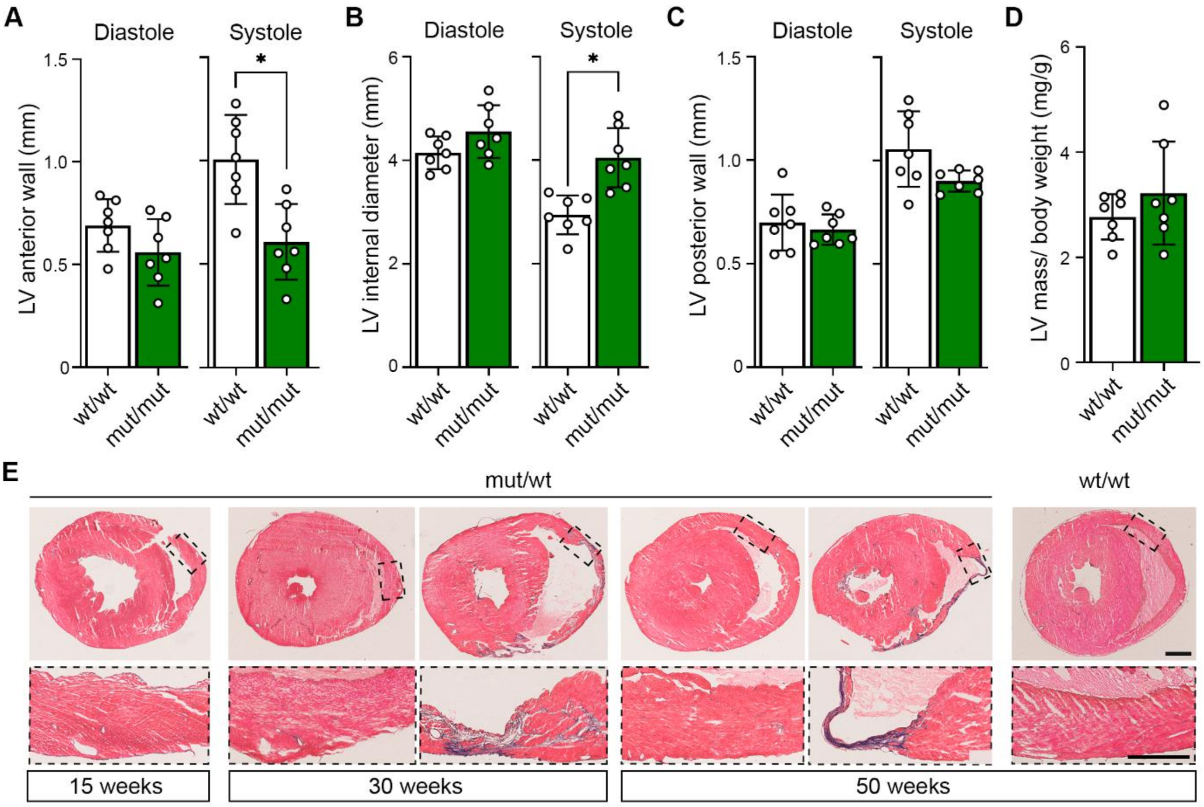


**Supplementary Figure 3. Additional data on DSG2-W2A mouse model.** (**A-C**) End-diastolic and – systolic measurements of left ventricular (LV) diameters by short axis (SAX) M-mode corresponding to Figure 2F - H. *P< 0.05, unpaired Student's t-test. (**D**) Calculation of the left ventricular (LV) mass by echocardiography. (**E**) Masson's trichrome fibrosis staining of DSG2-W2A mut/wt mice at the age of 15, 30 and 50 weeks corresponding to analysis in Figure 2K. Representative images of hearts with and without fibrosis are shown. Black rectangles depict area of zoomed insert. Scale bars: overview: 1 mm; insert: 0.5 mm.

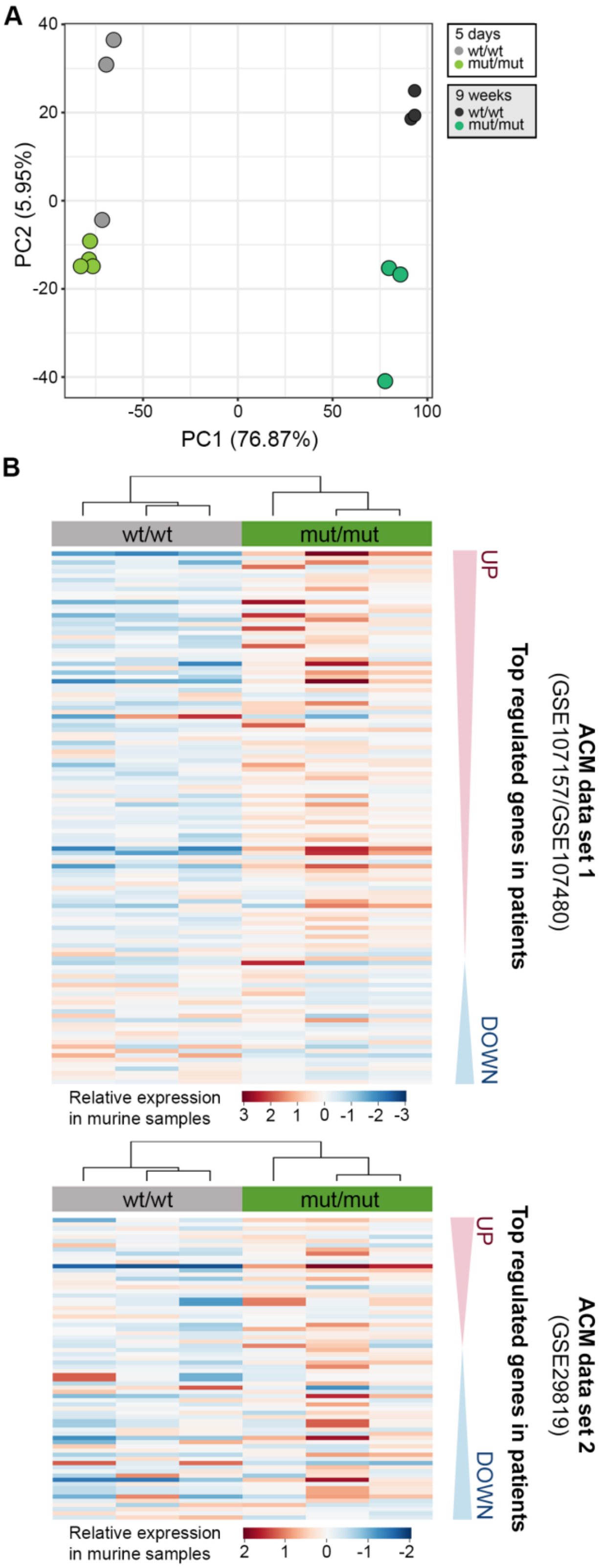


**Supplementary Figure 4. Validation of DGS2-W2A RNA sequencing data.** (**A**) Principal component analysis (PCA) of wt/wt and mut/mut hearts analysed by RNA sequencing at the age of 5 days and 9 weeks in two directories. (**B**) Heat maps of relative expression of the top differentially expressed genes derived from two ACM patient data sets (GEO: GSE107157/GSE107480 and GSE29819 [39]) in 9-weeks old mut/mut and wt/wt mouse hearts. Threshold of |Log fold change (FC)| > 2 and false discovery rate (FDR) < 0.05 was applied for gene selection. Samples are clustered based on their expression pattern. As depicted on the right, genes are arranged according to their expression in patient data sets with most upregulated genes on top to most down regulated on the bottom.

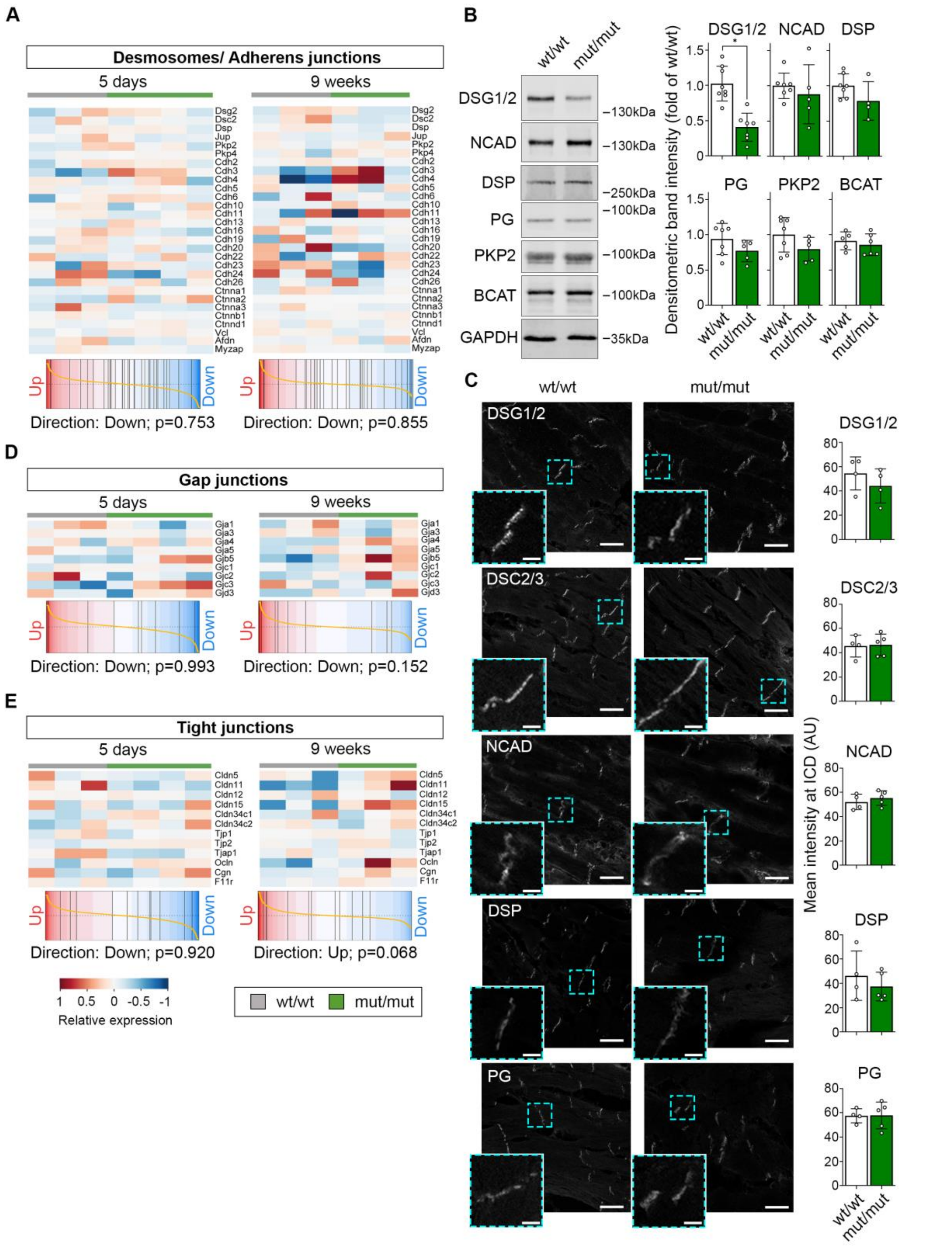


**Supplementary Figure 5. Junctional components are preserved in DSG2-W2A mutant hearts.** (**A**) Heat maps of relative expression of desmosomal and adherens junction components (genes indicated on the right) in 5-days and 9-weeks old mouse hearts. Gene set enrichment analyses of the included genes is depicted as barcode blot below. Samples are arranged according to their genotype (wt/wt: gray; mut/mut: green). (**B**) Western blot analysis and (**C**) immunostainings of adult DSG2-W2A wild type and mutant hearts with representative blots or images, respectively, on the left and related analysis on the right. Desmoglein-1/2 (DSG1/2), Desmocollin-2/3 (DSC2/3), N-cadherin (NCAD), desmoplakin (DSP), plakoglobin (PG), plakophilin-2 (PKP2) β-catenin (BCAT) were analysed as components of the mechanical junctions at the ICDs. GAPDH served as loading control in **B**. *P< 0.05, unpaired Student's t-test. Scale bars in **C**: overview 20 µm; insert 5 µm. Cyan rectangle depicts area of zoomed insert. (**D**, **E**) Heat map of relative expression of gap and tight junction molecules (genes indicated on the right). Gene set enrichment analyses of the respective genes is depicted as barcode blot below. Samples are arranged according to their genotype.

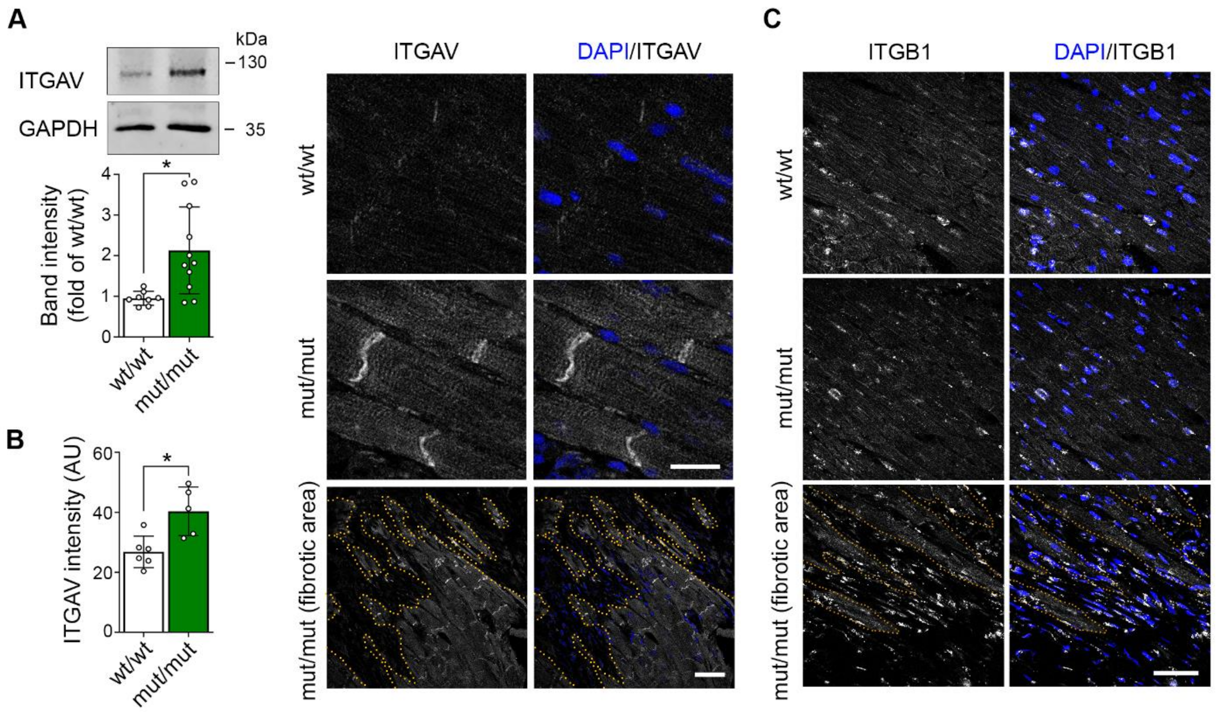


**Supplementary Figure 6. Integrin-αV is upregulated in DSG2-W2A mutant hearts.** (**A**) Western blot and analysis of band intensity of Integrin-αV (ITGAV) in DSG2-W2A hearts. GAPDH served as loading control. *P< 0.05, unpaired Student's t-test. (**B**) Immunostaining of ITGAV (red in overlay) in DSG2-W2A hearts. DAPI (blue) marks nuclei and F-actin (green) the sarcomere system. Lower row shows an overview image of a fibrotic area in mut/mut hearts. Dotted orange line marks the edge of fibrotic area. Scale bars: upper rows: 20 µm, lower row: 50 µm. (**C**) Integrin-β1 (ITGB1, white) immunostaining and nuclei counter stain (DAPI, blue) in DSG2-W2A mice. Lower row shows an overview image of a fibrotic area in mut/mut hearts. Dotted orange line highlights border of fibrotic tissue. Scale bar: 50 µm. Images representative for 5 mice per genotype.

| Gene name | Primer name | Sequence | Product size |
|---|---|---|---|
| *Col1a1* | qPCR_mCol1a1_for | CCCAGCCGCAAAGAGTCTAC | 152 |
| *Col1a1* | qPCR_mCol1a1_rev | GGACCCTTAGGCCATTGTGT | |
| *Lamc2* | qPCR_mLamc2_for | GTGCCGGAGTTACCATCCAA | 162 |
| *Lamc2* | qPCR_mLamc2_rev | CAGACATCAAGGGCCGAAGT | |
| *Fn1* | qPCR_mFn1_for | CTGGATCCCCTCCCAGAGAA | 193 |
| *Fn1* | qPCR_mFn1_rev | TTGGGGTGTGGAAGGGTAAC | |
| *Timp1* | qPCR_mTimp1_for | AGATACCATGATGGCCCCCT | 176 |
| *Timp1* | qPCR_mTimp1_rev | TGGTCTCGTTGATTTCTGGGG | |
| *Id2* | qPCR_mId2_for | ACATCAGCATCCTGTCCTTGC | 200 |
| *Id2* | qPCR_mId2_rev | ACGTGTTCTCCTGGTGAAATGG | |
| *Col3a1* | qPCR_mCol3a1_for | CCAGTGGCCATAATGGGGAA | 122 |
| *Col3a1* | qPCR_mCol3a1_rev | ATCTCGACCTGGCTGACCAT | |
| *Col1a2* | qPCR_mCol1a2_for | TGGATACGCGGACTCTGTTG | 87 |
| *Col1a2* | qPCR_mCol1a2_rev | GGCCCTTTCGTACTGATCCC | |
| *Itgb6* | qPCR_mItgb6_for | TGGCACTTCTGCCAAAGACT | 150 |
| *Itgb6* | qPCR_mItgb6_rev | TTTCTGTCTGGGCTCACGTC | |
| *Tubg2* | qPCR_mTubg2_for | GGTCTGGGCTCCTACCTCTTA | 96 |
| *Tubg2* | qPCR_mTubg2_rev | ACTCATCTCGTCCTGGTTGG | |
| *Gapdh* | qPCR_mGapdh_for | CCCACTCTTCCACCTTCGAT | 199 |
| *Gapdh* | qPCR_mGapdh_rev | AGTTGGGATAGGGCCTCTCTT | |

**Supplementary Table 1. Primers used in quantitative RT-PCR experiments.**
